# Adaptive Example Selection: Prototype-Based Explainability for Interpretable Mitosis Detection

**DOI:** 10.1101/2025.03.05.641711

**Authors:** Mita Banik, Ken Kreutz-Delgado, Ishan Mohanty, James B. Brown, Nidhi Singh

**Affiliations:** Pattern Computer, Inc., Friday Harbor, WA, United States; Department of Electrical and Computer Engineering, University of California San Diego, San Diego, CA, United States

## Abstract

Understanding the decision-making process of black-box neural network classifiers is crucial for their adoption in medical applications, including histopathology and cancer diagnostics. An approach of increasing interest is to clarify how the decisions of neural networks compare to, and perform parallel to, those of highly-trained and knowledgeable clinicians and other medical professionals within their prototypical classes of interest. Motivated by this, we introduce Adaptive Example Selection (AES), a prototype-based explainable AI (XAI) framework that facilitates the interpretability of deep learning models for mitosis detection. AES works by selecting and presenting a small set of real-world mitotic images most informative to a given classification, allowing pathologists to visually assess and understand the neural network’s decision by comparing test cases with similar previously annotated examples. AES achieves this by expanding the neural network’s confidence/belief function and fitting it to a radial basis function (RBF) approximator, an approach we term Decision Boundary-based Analysis (DBA). This method makes the decision boundary more transparent, offering robust visual insights into the model’s decisions, and thereby equipping clinicians with the information needed to effectively utilize AI-driven diagnostics. Additionally, AES includes customizable user controls, allowing clinicians to tailor decision thresholds and select prototype examples to better align with their specific diagnostic needs. This flexibility empowers users to engage with the AI model more directly and meaningfully, increasing its practical relevance in clinical settings.

## 1 Introduction

Mitosis, the process of tumor cell division, serves as a critical biomarker in many human and animal cancers, with histopathological analysis remaining the gold standard for cancer diagnostics Gurcan et al. (2009). Traditionally, pathologists manually detect and count mitotic figures—dividing cells—within regions exhibiting the highest proliferative activity. However, this process is time-consuming, highly variable, and heavily reliant on the pathologist’s expertise. Several challenges make mitosis detection difficult. The four phases of mitosis—prophase, metaphase, anaphase, and telophase—exhibit different nuclear shapes and textures, complicating identification. Additionally, non-mitotic cells such as lymphocytes and apoptotic cells, may resemble mitotic figures in hematoxylin and eosin–stained (H&E–stained) images, increasing false positives. Pathologists may also miss small or focally present mitotic figures, especially when reviewing numerous slides under time constraints, leading to subjectivity and variability Balkenhol et al. (2019). These factors, along with inconsistencies between and within individual raters, negatively impact diagnostic accuracy. However, it is important to note that no existing AI approach has achieved super-human performance in this high-trust task – nor is that our goal. Instead, we aim to augment classification accuracy, particularly in challenging cases that fall close to human or machine decision boundaries European Society of Radiology (ESR) (2022).

Recent advancements in deep learning (DL) and digital pathology have sparked increasing interest in automating mitosis detection Veta et al. (2016); Bertram et al. (2022); Aubreville et al. (2023a), offering the potential to enhance workflow efficiency and reduce the intra- and inter-rater variability. van der Laak et al. (2021). DL models typically analyze high-resolution, digitized whole slide images (WSIs) of H&E-stained tissues, which can contain tens of millions of pixels and are typically annotated by pathologists. These algorithms identify mitotic figures by placing bounding boxes around cells of interest. Some research groups have demonstrated DL performance comparable to human pathologists Aubreville et al. (2020). However, despite increasing powerful and generalizability, DL models are often criticized as inscrutable “black boxes” due to their highly complex, multi-layered neural networks with millions of parameters. This lack of transparency poses a significant barrier to clinical adoption Jia et al. (2020), particularly in medical applications where the consequences of errors can be far-reaching.

In gaining clinical acceptance, AI models must incorporate explainability. The ultimate source of information for patients is the clinician, a relationship that cannot be replaced by automation. To effectively pass on insights derived in part from automation, clinicians must trust the models as much as their patients trust them Murdoch et al. (2019). *Explainable AI* (XAI) allows practitioners not only to understand how a model arrives at its predictions but also to assess its confidence levels, limitations, and potential biases. Effective XAI systems should allow for active user feedback, enabling pathologists to adjust model sensitivity, fine-tune performance, and provide feedback to build trust in AI-assisted diagnostics. As explainability needs varies across users and domains, it is essential to evaluate these systems through iterative user studies to ensure XAI explanations are clear, relevant, and actionable Evans et al. (2022); Plass et al. (2023).

In histopathology, several XAI techniques have been developed to make DL models more explainable. These fall into two main categories: heat maps (or saliency maps) Selvaraju et al. (2017); Ribeiro et al. (2016); Lundberg and Lee (2017) and example-based methods Hoffer and Ailon (2015); Koh and Liang (2017); Chen et al. (2019). Saliency maps highlight the areas of an image that contributed most to a model’s decision, giving users a visual sense of the decision-making process. However, they often lack precision Rudin (2019), as they highlight broad regions rather than specific cellular features such as mitotic figures, cancerous cells, or other microscopic structures. This ambiguity makes it difficult for pathologists to interpret the model’s focus, reducing trust and limiting their usefulness in clinical decision-making. Conversely, example-based reasoning refers to systems that provide insights into decisions about a test case (e.g., a new medical image) by contrasting it with previous cases that the model has encountered and classified Hegde et al. (2019). Such methods provide a clear rationale for the model’s decision by showcasing reference cases, making it easier for clinicians to interpret the results. This approach aligns with the core principles of example-based reasoning—leveraging prior knowledge to reason through new problems.

Prototype-based approaches, a subset of example-based reasoning, aim to explain data embeddings, and therefore decision boundaries, by displaying clinically-relevant real-world examples indicative of a particular classification decision Chen et al. (2019); Li et al. (2017). These explanations reference a set of contextually representative prototypical cases that the model uses to justify its predictions. Despite the promise of prototype-based methods, few studies have successfully integrated their algorithmic implementation with domain-specific, user-centric requirements—a necessary step to make XAI tools more intuitive and relatable in clinical settings.

Here, we introduce Adaptive Example Selection (AES)—a prototype-based XAI framework designed to clarify the decision-making process of deep learning models for mitosis detection, aiming to build end-user trust and confidence in adopting AI solutions in digital pathology. Figure 1 illustrates an end-to-end XAI-based mitosis detection workflow for cancer screening. Our approach consists of two key steps: (a) training a black-box Faster R-CNN Mitotic Figure (MF) detector to identify mitotic cells in histopathology images, and (b) developing the AES probing mechanism to analyze and visually explain the detector’s confidence scores, allowing for interactive interpretation of bounding box predictions. By integrating AES, we aim to make DL-based mitosis detection more intuitive and trustworthy for pathologists, bridging the gap between AI development and clinical usability.

**Figure 1:**
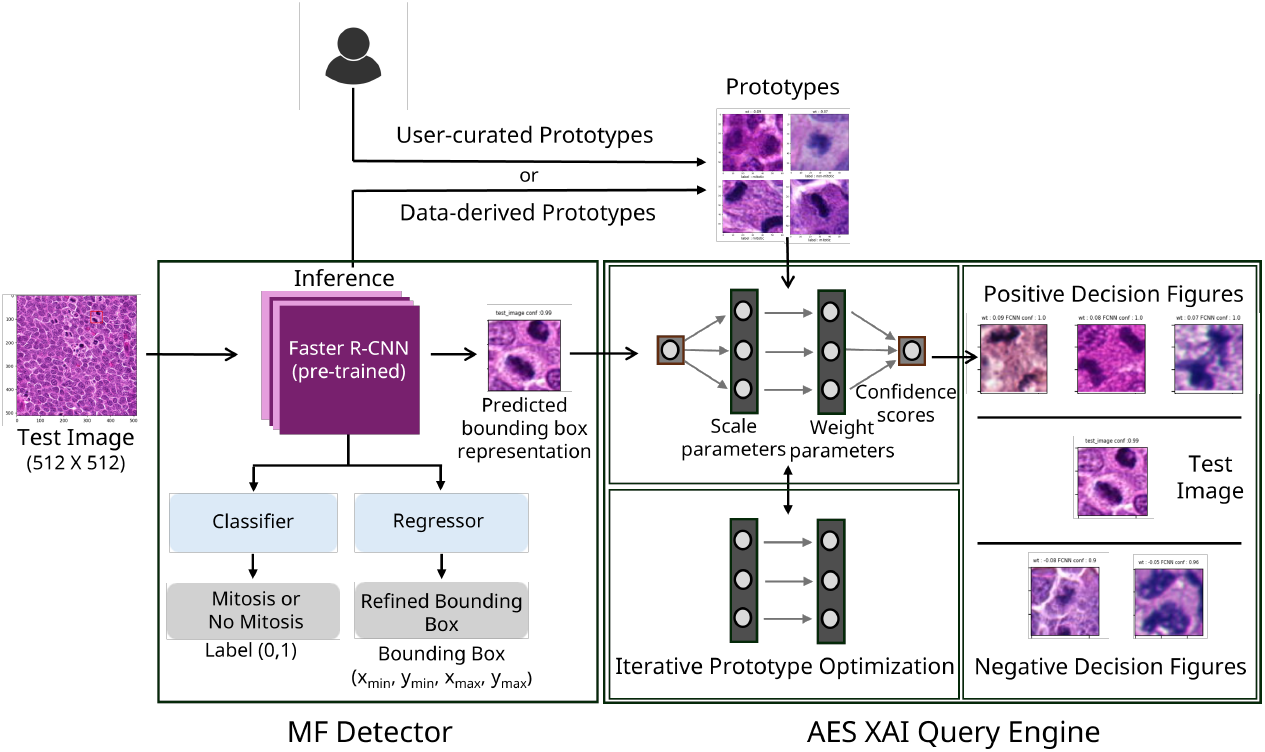
Overview of the XAI-driven mitosis detection workflow for cancer screening. The Faster R-CNN Mitotic Figure (MF) Detector identifies localized bounding boxes of interest in histopathology images provided to the detector. The Adaptive Example Selection (AES) module provides interpretability by retrieving prototypical example bounding box images that correspond to expert-labeled mitotic and nonmitotic figures, thereby providing meaningful visual explanations.

## 2 Results

In this section, we present the outcomes of our experiments evaluating the effectiveness of the AES framework in enhancing the interpretability of mitosis detection in histopathology images. We begin by assessing the performance of the MF Detector, focusing on its classification accuracy and detection reliability. We then evaluate the AES query engine’s ability to generate meaningful visual explanations by retrieving prototypical examples that align with expert-labeled mitotic and non-mitotic figures.

### 2.1 Faster R-CNN Mitotic-Figure Detector: Training and Validation

In the initial phase of our experiment, we train and deploy a region of Interest (ROI)-based DL model to detect mitotic figures in histopathology images. This is a binary classification problem: a positive identification indicates that a mitotic figure has been correctly localized, while a negative identification means the detected object is not a mitotic figure. The model identifies potential objects of interest by placing a bounding box (ROI) around them and assigning a confidence score—a learned function value that represents the model’s belief that the detected object is a mitotic figure.

Our model utilizes a two-stage Faster R-CNN architecture Ren et al. (2015), trained on the publicly available MIDOG++ dataset, denoted as ℳ, which has been labeled by trained clinicians as described in Aubreville et al. (2023b). Using training data ℳ_train_ ⊂ ℳ extracted from ℳ, our model first generates a region proposal (a bounding box, represented by a vector *x* of parameter values, of an object detected in an image). It then refines the proposal using the confidence score provided by a learned belief function *β*(*x*) ∈ [0, 1] where *β*(*x*) represents the “confidence” that the bounding box contains an object of positive type (mitotic figure).

A detection is classified as a mitotic figure if *β*(*x*) ≥ *τ*_0_, where *τ*_0_ ∈ [0, 1] is a user-defined threshold. In our experiments, we set *τ*_0_ = 0.969 to optimize the F1-score through cross-validation on a hold-out test set ℳ_test_ ⊂ ℳ, ensuring that ℳ_test_ ∩ ℳ_train_ = ∅. We evaluated model performance using three key metrics: precision, recall, and F1-score, as summarized in Table 1. For reference, we also include the F1 scores reported in Aubreville et al. (2023b). These metrics offer a comprehensive assessment of the model’s effectiveness, with the F1-score summarizing the balance between precision and recall.

**Table 1:** Summary of the MF Detector’s performance across different tumor types. ^†^For comparison, F1-scores from Aubreville et al. (2023b) are also included.

| **Tumor Type** | **Precision** | **Recall** | **F1-score** | **F1-score<sup>†</sup>** |
| --- | --- | --- | --- | --- |
| Human Breast Cancer | 0.79 | 0.73 | 0.76 | $0.71 \pm 0.02$ |
| Canine Lung Cancer | 0.64 | 0.72 | 0.68 | $0.68 \pm 0.02$ |
| Canine Lymphosarcoma | 0.72 | 0.81 | 0.76 | $0.73 \pm 0.01$ |
| Canine Cutaneous Mast Cell Tumor | 0.80 | 0.91 | 0.85 | $0.82 \pm 0.01$ |
| Human Neuroendocrine Tumor | 0.63 | 0.51 | 0.57 | $0.59 \pm 0.01$ |
| Canine Soft Tissue Sarcoma | 0.77 | 0.75 | 0.76 | $0.69 \pm 0.01$ |
| Human Melanoma | 0.78 | 0.78 | 0.78 | $0.81 \pm 0.01$ |

### 2.2 AES Query Engine: Training, and Validation

Having trained a black-box MF detector, we developed AES, a visually intuitive prototype-based XAI interrogation engine, to analyze and explain the MF detector’s classification decisions in a manner that is consistent with clinician training practices, and hence familiar to the end user.

#### 2.2.1 Glossary of Notations

This subsection provides a comprehensive overview of the notations used throughout the article, specifically in the context of bounding boxes derived from trained MF detector and the AES query engine. Readers are encouraged to refer back to this section as needed for clarity on terminology and symbols.

- *x* = parameter values that localize and give the shape of a rectangular bounding box.
- ℬ ^*all*^ = set of all bounding boxes *x* found by the ROI object detection algorithm.
- The Faster R-CNN algorithm learns a belief function *β*(·) that maps to confidence scores *β*(*x*) ∈ [0, 1] used to classify a box *x* ∈ ℬ ^*all*^ as being a mitotic figure.
- A MF detection threshold value *τ*_0_ ∈ [0, 1] is selected and used to identify a set of *τ*_0_–level positively predicted bounding boxes,

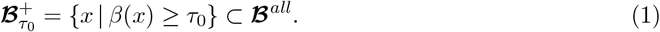
- We define additional subsets of ℬ ^*all*^ according to the general definition

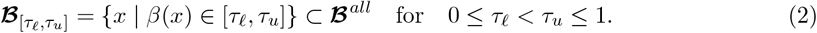
- We define a subset ℐ ⊂ ℬ ^*all*^ of *figures of interest* identified by a subject matter expert as being worthy of paying special attention to.
- Given *τ*_0_ and ℐ we define the subsets PDF and NDF of *positive and negative decision figures at margin ϵ* ≥ 0 respectively by

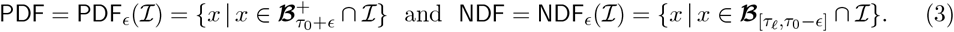
- *r*_*x*_ = latent variable representation of bounding box *x* ∈ ℬ ^*all*^ provided by a neural network. 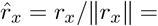 normalized latent variable representation of bounding box *x*.
- *i*_*x*_ = image contained in bounding box *x* ∈ ℬ ^*all*^.
- We write *i*_*x*_ ≐ MF if the image is that of a mitotic figure and *i*_*x*_ ≐ NMF otherwise.
- Unless otherwise indicated, the quantities defined above hold for the training data set ℳ_train_. When defined for a test data set, this is indicated as 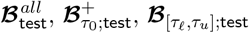 etc.

For succinctness we will typically refer to *x*∈ ℬ^*all*^ as “the bounding box *x*”; *r*_*x*_ as “the latent variable for *x*”; 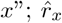 as “the normalized latent variable for *x*”; and *i*_*x*_ as “the image or prototype for *x*.” Furthermore, when the context should be clear, we may at times variously refer to *x* as “the” bounding box, representation, normalized representation, or image of interest.

#### 2.2.2 Transforming Beliefs for Enhanced Decision Boundary–based Analysis

Our AES query engine aims to facilitate an expanded view of decision making for predicted bounding boxes *x* near, and relative to, the decision threshold value *τ*_0_. This is done by transforming the belief/confidence scores *β*(*x*) as follows: First, they are transformed using a variance-stabilizing Box-Cox(BC) transformation; Second, they are then centered at the transformed value of the detection threshold *τ*_0_. This results in centered BC transformed confidence scores *β*(*x*) → *f* (*x*) according to the isomorphic mapping^2^

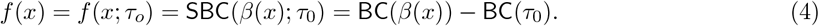

The sets of *ϵ*-margin positive and negative decision figures PDF = PDF_*ϵ*_(I) and NDF = NDF_*ϵ*_(I) can be written in terms of decision thresholds placed on the transformed belief function *f* (*x*). The procedure of “zooming-in” followed by a fine-grained analysis of classified images relative to a specified decision threshold *τ*_0_ we call (enhanced) *Decision Boundary Analysis* (DBA). A typical result of the expansion *β*(*x*) → *f* (*x*) is shown in the lower left-hand corner of Figure 2.

**Figure 2:**
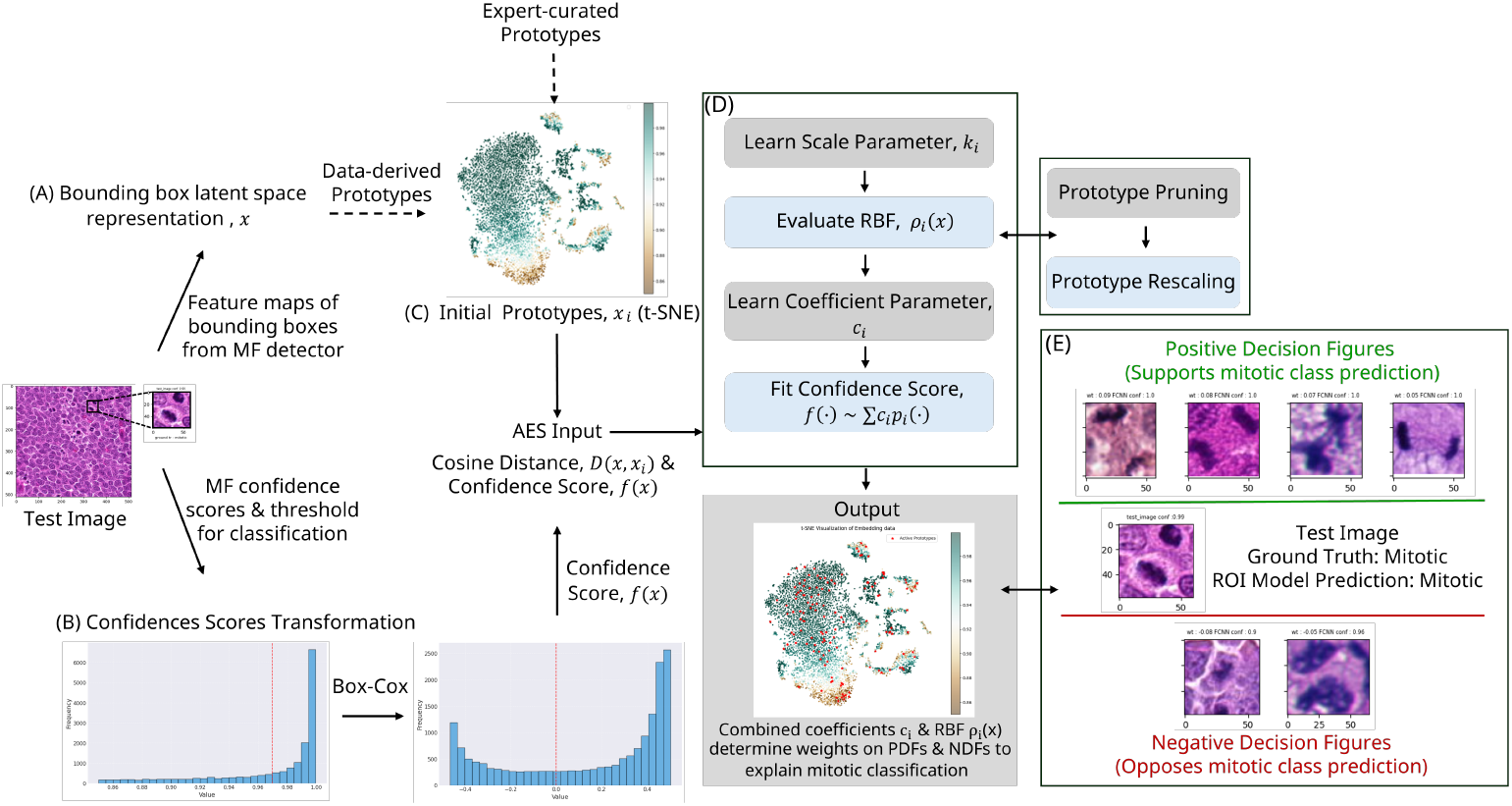
The Adaptive Example Selection (AES) Engine: (A) Feature maps and corresponding confidence scores for MF classification are extracted for each predicted bounding box in a test image. (B) Confidence scores are normalized using a variance-stabilizing power transformation to enhance the MF detector’s behavior at the decision boundary. (C) Initial prototypes of mitotic and non-mitotic figures are selected either from the training data or an expert-curated database. (D) During the AES optimization training stage, prototypes are pruned, and RBFs are scaled to select a sparse, informative dictionary of active prototypes for inference. (E) During inference, for each predicted bounding box, the trained AES assigns coefficients that quantify the influence of nearby prototypical PDF (Positive Decision) and NDF (Negative Decision) figures (mitotic and non-mitotic), providing visual insights into mitotic class predictions.

#### 2.2.3 DBA and Approximation of the Transformed Classifier Belief Function

Rather than focusing on the (necessarily discontinuous) binary decision itself, our DBA procedure instead models the transformed belief function *f* (·) via an approximation, 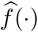, determined from a radial basis function (RBF) regression. Each term (say the *i*-th term) of the regression 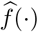 is centered on an example clinical image 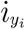 of known label (positive or negative) provided by a subject matter expert, and an appropriately chosen collection of such images associated with a value 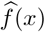 of the approximation for the belief function *f* (*x*) (equivalently, for *β*(*x*) ↔*f* (*x*)) can be provided to a clinician to gain insight into what labelled actual example images inform that degree of belief that *i*_*x*_ is a mitotic figure.

We use localized (at *y*_*i*_) and trimmed (symmetrically truncated support) RBFs 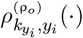 (see below) to approximate the transformed confidence score function via

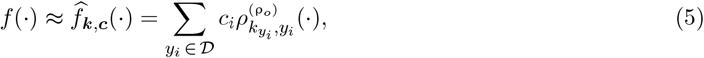

where 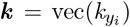 and ***c*** = 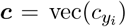 are respectively vectors with elements 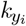 and 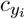 for localization (i.e., RBF center) points *y*_*i*_ ∈ 𝒟 ⊂ 𝒟 _0_, where 𝒟 _0_ ⊂ ℬ ^*all*^ is an initializing set of bounding boxes in ℬ ^*all*^ that are *known-label* actual example images (exemplary actual images of *both* mitotic and non-mitotic cells) and 𝒟 is a post-training set of the bounding boxes of learned prototypical actual example images. We also set

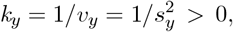

where *s*_*y*_ *>* 0 functions as a *y*-dependent scale factor and, similarly to ***k***, we define the vectors ***v*** and ***s*** in a component-wise manner.^3^

We call the parameter *k*_*y*_ the *concentration* of the RBF centered at 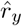, the parameter *v*_*y*_ the *variation* of the RBF centered at 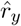, and the parameter *s*_*y*_ the *scale* of the RBF centered at 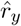. We have defined the concentration vector ***k*** to be the vector of concentration values *k*_*y*_ for all values of *y* ∈ 𝒟 ⊂ 𝒟_0_, where we are interested in an initially selected set of *admissible* RBF centers,

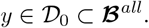

The elements of 𝒟_0_ are known to be the bounding boxes of representative actual (mitotic and non-mitotic cell) figures which are selected by a subject matter expert.

For a specified truncation parameter ρ_0_ each truncated RBFs in (5) is of the form

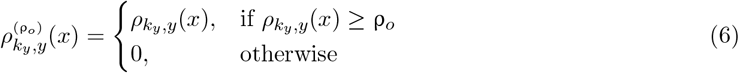

with

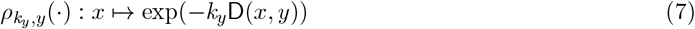

where

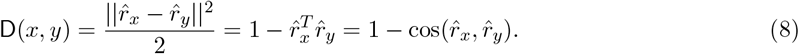

The function 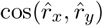 is the cosine similarity between the normalized representations 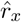 and 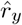 Note that the functions (6) and (7) are not symmetric with respect to the interchange of *x* and *y*, which is unlike the case encountered in Kernel Method-based classification. In our experiments we set the trim parameter to *ρ*_0_ = 0.1.

#### 2.2.4 Training and AES Optimization Framework

The fit of 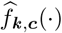 to *f* (·) is attained by performing the functional approximation optimization corresponding to alternating between Equations (9) and (10). The (constrained) optimization is done with the goal of learning a small and representative collection 𝒟 ⊂ 𝒟_0_ of explanatory prototypical examples that are useful for understanding the decision making behavior of the MF detector and classification algorithm. The final, adaptively learned, set 𝒟 provides a dictionary of prototypical examples which is usefully bounded in size (specifically, the size of | 𝒟| is kept from becoming redundantly or uninformatively large) while containing highly informative features used to determine a succinct set of images that provide a visual explanation of predictions made about new data.

Towards this end, we perform the following two optimizations to learn the prototypical examples belonging to 𝒟, a procedure which we refer to as AES. This is done by alternatively minimizing the two shown loss functions ℒ _1_(***c, k***) and ℒ _2_(***k***) with respect to their arguments and the constraints shown below:^4^

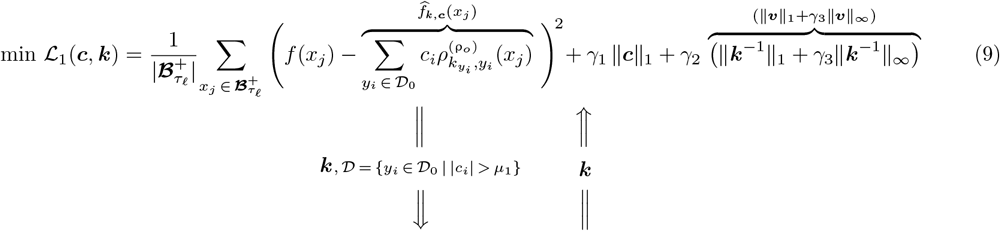

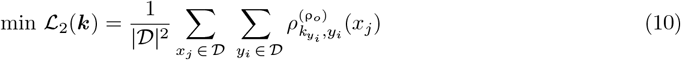

subject to the sign-alignment constraints

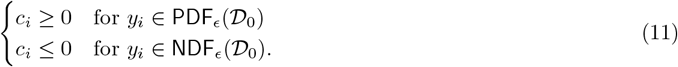

At convergence, the optimization yields a dictionary of prototypical examples

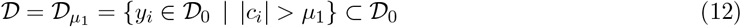

where the parameter *µ*_1_ *>* 0 sets the minimum size for a coefficient *c*_*i*_ to be considered to be relevent, a concentration vector 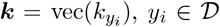, and a coefficient vector ***c*** = vec(*c*_*i*_), for *i* such that *y*_*i*_ ∈ 𝒟. Note that the effective dimensionality for both of the vectors ***k*** and ***c*** is equal to | 𝒟 |, the size of 𝒟. We similarly define vectors ***y, v***, and ***s***.

Once the optimization phase of the AES procedure is completed, an approximation of *f* (*x*) over the learned set of global dictionary of prototypical examples 𝒟 is provided by Eq. (5). Further, from 𝒟 a local image-dependent set AES(*x*) of prototypical examples associated with a new image *i*_*x*_ (contained in the ROI R-CNN-determined bounding box *x*) is extracted. The local set AES(*x*) is defined by

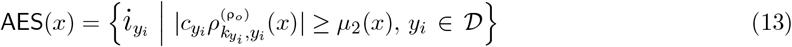

for a parameter *µ*_2_(*x*) *>* 0 that is equivalent to taking AES(*x*) be the set of active prototypes for *x* that explain a significant percent (for example, 90%) of the prediction as discussed in the Methods Section 5.2.5.

The AES methodology queries the decision of the MF detector by allowing a subject matter expert (e.g., clinician or researcher) to gain explanatory insight. This is achieved by comparing the classification decision for a new image *i*_*x*_, with prototypical example images and their known classification labels in the set AES(*x*). Examples of this procedure are shown in Figures 2.

#### 2.2.5 Principles of Enhancing Explainabilty Embodied in the AES Methodology

Our XAI-motivated adaptive selection of prototypical examples (see the workflow in Figure 2) follows well-established principles of feature interpretability, including *localization* (of latent space representations), feature *sparsity*, and *non-negativity*. More broadly, our approach employs *sign-alignment* to ensure that features used for interpretability correspond with the decisions made by the MF detector. These properties are interrelated (not necessarily independent) and aspects of our optimization strategy influence multiple properties simultaneously.

Localization, distinctiveness, and sparse selection of prototypical features are ensured by locating finite-support (parameterized by) 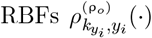 on separate prototypical images *y*_*i*_. These images are trained to be sparsely selected (via the use of the 1-norm regularization in the ℒ _1_ loss function minimization of Eq. (9)), and spread out and well-isolated from each other^5^ when selecting *y*_*i*_-dependent concentration parameters 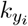 via the minimization of the loss function ℒ _2_ in (10). Note that bringing in the ℒ _2_ optimization provides an *iterative refinement step* that enhances the distinctiveness of prototypical examples beyond the standard sparsity-enforcing 1-norm optimization of Eq. (9).

Since our goal is to interrogate MF detector’s decisions, we impose the sign-alignment constraints (11) to ensure that training images that are above the decision threshold of the detector remain above the threshold in our approximation Eq. (5), and vice versa.^6^ By enforcing consistency with the detector’s decisions, especially near the threshold-decision region, we can carefully scrutinize which images lead to incorrect classifications, revealing potential sources of “confusion” in the detector’s decision-making process.

#### 2.2.6 Validation of the AES XAI Methodology

The learned dictionary 𝒟 provides a *globally* sparse representation of the training data set 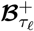. If the training data set is sufficiently representative and our training methodology is robust, this representation should also generalize to any properly sampled test dataset 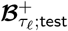. Conceptually, all data samples (both training and text) are drawn from a data manifold for which the sparse (in the sense that 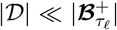) set 𝒟 can be use to accurately represent a decision about any specific (localized) point 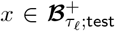 via the use of the *localized* sparse approximation (See Eq. (17) in the Methodology section) which depends on the localized sparse approximation set AES(*x*), where generally 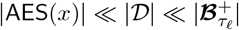. The elements in the globally sparse dictionary 𝒟 serve as candidate explanatory prototypical examples for explaining a decision made about any new datum *x* while the elements in the locally sparse set AES(*x*) are the actual “localized” explanatory prototypes used to explain the decision.

After convergence of the approximation procedure, we assess the quality of the RBF fit (5)using the learned RBF concentration parameters ***k*** and RBF centers (the learned prototypical examples) 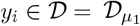 by evaluating the condition number *κ*(D) of the RBF matrix^7^

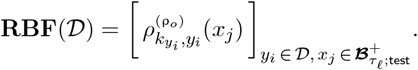

The condition number *κ*(𝒟) quantifies how well the sparse set 𝒟 is “spread out” on the underlying data manifold that any data set 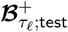 is presumed to be sampled from, with a lower value of *κ*(𝒟) corresponding to better coverage by 𝒟 of the data set 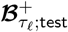. Our numerical experiments demonstrate that incorporating variable (*y*-dependent) concentration parameters *k*_*y*_, along with truncated RBFs and and the use of the alternate optimization (9)–(10), significantly improves the conditioning of the matrix **RBF**(𝒟). This enhancement leads to more informative prototypes, consistent with prior research Kansa (1990); Kansa and Carlson (1992) that shows that using location-dependent scale parameters 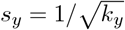 improves performance of RBF-based functional approximators.

To evaluate the effectiveness of the AES query engine, we conducted extensive experiments using metrics that assess its ability to: (a) achieve global sparsity, determined by the number | 𝒟| of candidate active prototypes contained in the dictionary | 𝒟 |, (b) approximate confidence scores via the use of the *localized* sparse approximation (17), as measured by *R* values, and (c) minimize the number of relevant active prototypes per prediction of new observations *x*,referred to as relevance scores (RS) measured by the mean (denoted by Mean-RS) and median (denoted by Med-RS) of the size |AES(*x*)| of the actual explanatory sets AES(*x*), subject to the constraint that the approximation (17) explains over 90% of the prediction. Key results from these experiments are summarized in Table 2.^8^

**Table 2:** Performance of the AES interrogation algorithm with different forms of the approximation model (5) and training optimization strategies (9)–(10). “No trim” (ρ_0_ = 0 in RBF 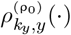) means that support truncation of the RBF has been turned off. Setting ρ_0_ = 0.1 means that the RBF supports are truncated in the manner described in Eq. (6) and Eq. (S19)–(S21). “Variable ***k*** “ means each component of ***k*** can be different and is learned accordingly during training. “Constant *k*” means all components of ***k*** take the same value *k*, which is learned during training. “ℒ_2_ off” means only the optimization of Eq. (9) is performed during training. “ℒ_2_ on” means the alternating optimization (9)–(10) is performed. Parameters *γ*_1_ = 0 and/or *γ*_2_ = 0 means the corresponding regularization terms in optimization (9) are turned off.

| Row | Case Study | Parameter Values | $R^2$ | Mean-RS | Med-RS | $\vert \mathcal{D}\vert $ | $\log_{10}(\kappa(\mathcal{D}))$ |
| --- | --- | --- | --- | --- | --- | --- | --- |
| 1 | $\rho_{k_y, y}^{(0)}$ no trim; $\mathcal{L}_2$ off | constant $k$, $\gamma_1 = \gamma_2 = 0$ | 0.97 | 606.95 | 146.10 | 1647.50 | 5.70 |
| 2 | | constant $k$, non-zero $\gamma_1, \gamma_2$ | 0.95 | 253.94 | 238.90 | 1576.20 | 4.93 |
| 3 | | variable $\mathbf{k}$, non-zero $\gamma_1, \gamma_2$ | 0.96 | 268.85 | 245.90 | 620.90 | 4.46 |
| 4 | $\rho_{k_y, y}^{(0.1)}$ trimmed; $\mathcal{L}_2$ off | constant $k$, $\gamma_1 = \gamma_2 = 0$ | 0.96 | 621.50 | 791.00 | 1619.10 | 4.16 |
| 5 | | constant $k$, non-zero $\gamma_1, \gamma_2$ | 0.94 | 182.03 | 210.60 | 1844.50 | 4.36 |
| 6 | | variable $\mathbf{k}$, non-zero $\gamma_1, \gamma_2$ | 0.96 | 21.17 | 8.50 | 269.40 | 2.97 |
| 7 | $\rho_{k_y, y}^{(0.1)}$ trimmed; $\mathcal{L}_2$ on | variable $\mathbf{k}$, non-zero $\gamma_1, \gamma_2$ | 0.96 | 14.81 | 10.00 | 190.20 | 2.87 |

In Table 2, the *R*^2^–column indicates the quality of the fit

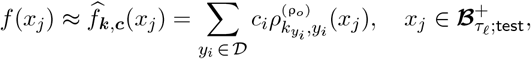

averaged over the test data samples 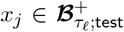 for the shown various training parameter values. The | 𝒟|–column gives the size of the learned global dictionary of explanatory prototypical examples 𝒟. The Mean-RS–column and Med-RS–column are measures of the average value of the size |AES(*x*)| of the local dictionary of explanatory prototypical examples AES(*x*) ⊂ 𝒟.

When *γ*_1_ and *γ*_2_ are nonzero, the sparsity-enforcing regularization terms in the loss function ℒ_1_ of Eq. (9) become active.^9^ This ℒ_1_ loss minimization is similar to Kernel-Lasso (in the RKHS setting, Roth (2004)) or RBF-Lasso in the general setting, both of which generate sparse coefficients. We employ variable scale parameters (equivalently, variable concentration parameters ***k***) as they intuitively adapt to regions with varying data distributions Bozzini et al. (2003). This leads to improved localization, fewer global prototypes, a reduced number of relevant examples per prediction, and lower condition numbers.

Note that some cases impose constraints of constant scale: for all 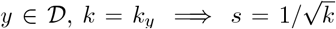. When *k* is constant and *γ*_1_ = *γ*_2_ = 0, the optimization of ℒ_1_ of Eq. (9) corresponds to a standard quadratic loss function optimization.^10^

The first six rows of Table 2 are trained ignoring the optimization of the loss ℒ_2_ shown in Eq. (10). The role of the ℒ_2_ minimization in the full alternating minimization procedure (9)–(13) is to enhance distinctiveness of the RBFs by enforcing concentration of each RBF independently of their collective action as regression basis functions, which is being forced via the ℒ_1_-optimization of Eq. (9)). Loss functions ℒ_1_ and ℒ_2_ represent competing goals and their combined optimization falls under the rubric of the theory of multi-objective optimization (see Table S3 in the Supplementary section). To address this, we use a heuristic approach with early stopping in the ℒ_2_ optimization. This creates a new initial condition of the value of ***k*** for the subsequent cycle of ℒ_1_ optimization, attempting to push the ℒ_1_ optimization into a more fruitful basin of attraction. Our experiments confirm that this does indeed appear to be the case.

Rows 1-3 of Table 2 correspond to the use of non-truncated RBFs,

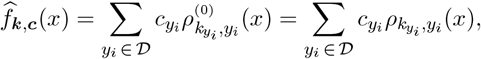

with the RBF distinctiveness optimization (10) turned off. We then create a simple baseline model based on taking *k* = constant (constant scale RBFs) and turning off sparsity regularization by setting *γ*_1_ = *γ*_2_ = 0. Since in this case the loss function is purely focused on quadratic loss minimization of the functional approximator, we obtain the best *R*^2^ value. However, this comes at the expense of the largest global dictionary size | 𝒟| and the largest average size of the local dictionary |AES(*x*)|. Furthermore, ignoring regularization in the optimization results in the largest value of the condition number *κ*(𝒟), which suggests possible over-fitting to the data. Interestingly, for the case of non-truncated RBFs, when regularization is turned on (non-zero *γ*_1_ and *γ*_2_) we see significant improvement in the sparsity (relative to the size of the baseline) of the global dictionary and robustness as measured by the condition number for the case of non-constant concentration but slight degradation in the average size of |AES(*x*)|.

Rows 4-6 of Table 2 reproduce the conditions of rows 1-3 but with truncated RBFs to enhance compactness and distinctiveness. Here, improvements are unambiguous only for the case of a nonconstant concentration vector ***k*** (row 6), where performance dramatically improves across all measured criteria.

Building on row 6, we introduce ℒ_2_ optimization to further enhance the placement and separation of RBFs, leading to improved performance. As shown in row 7, this enhancement improves three of the four performance metrics, with a significant improvement in the size of the global dictionary | 𝒟 |.

The intuition behind using trimmed (truncated) RBFs is to achieve compact support of the basis functions. The compact support aims to minimize the influence of distant points during local interpolation. This enhances stability and interpretability during inference. While smooth compactly supported radial functions like Wendland functions Wendland (1995) could be used instead, we opted for discontinuously trimmed (i.e., truncated) Gaussian functions due to their straightforward implementation. As shown in Table 2, using compactly supported basis functions leads to a further reduction in relevance scores and condition numbers, which comes with a slight trade-off in performance, as indicated by the *R*^2^ values.

Given the insights from rows 1-6 of Table 2, it is natural to attempt to further encourage distinct placement and separation of the RBFs. Toward this end, we implemented an alternating optimization approach that alternates between the competing loss functions ℒ_1_ and ℒ_2_. This approach aims to improve the distinctness of the pruned/active prototypes by adjusting the concentration parameters of the RBFs via the incorporation of the optimization shown in Eq. (10). We numerically observe that this approach lowers mean relevance scores and condition numbers with minimal performance degradation. By concentrating weight and coefficients on fewer active prototypes, it minimizes redundancy during inference. Concentrating weight on fewer chosen prototypes has also been explored in *ProtoFac*, Das et al. (2020) and *ProtoAttend* Arık and Pfister (2020). However, while alternate optimization considerably lowers mean relevance scores, it does not produce a corresponding reduction in median relevance scores.

## 3 Discussion

In this work, we introduce Adaptive Example Selection—a prototype-based query engine—designed to explain the decisions of a DL-based model for mitotic figure detection. For a given detection decision made by the model, AES retrieves real-world prototypical example images that inform decision. These examples, drawn from a curated clinical database of mitotic-figure images annotated by subject matter experts, help end-users understand and trust the Faster R-CNN model’s outputs. Developing user-centric explainability for black-box DL models is crucial for their adoption in high-stakes environments like pathology, where clinicians must trust and understand AI-driven decisions due to their significant impact on patient outcomes. Without addressing the explainability gap, these models remain opaque, leading to skepticism and hindering their widespread use. By creating explanations that align with the mental models and expertise of pathologists, AES helps bridge this trust gap, promoting AI integration into diagnostic workflows.

The core challenge in designing XAI systems is determining what type of explanations users understand and trust, especially in the absence of standardized explainability metrics. Published literature on state-of-the-art explainability techniques in digital pathology Chen et al. (2024); Evans et al. (2022) has shown that prototype-based explanations are among the most desirable. Prototypical examples are less disruptive to users’ intuition, encourage inductive reasoning about features and decision tasks, and provide strong, reliable signals when predictions are uncertain. Building on these insights, we developed AES to enhance explainability by comparing test images with real clinical examples (rather than synthetically generated prototypes). These prototypes are sourced from either the training dataset or user-provided examples, allowing users to better interpret the model’s results and reasoning.

### 3.1 Faster R-CNN Mitosis Figure Detector

The first step in our approach was developing a region-of-interest (ROI) detection model for identifying mitotic figures in histopathology images. We selected the Faster R-CNN architecture (Faster R-CNN; Ren et al. (2015)) for its superior accuracy over one-stage detectors like YOLO Cai and Vasconcelos (2022), particularly for complex biomedical imaging tasks. To enhance performance, we integrated a ResNet-50-FPN backbone, which improved both classification accuracy and spatial localization. Additionally, we incorporated a Feature Pyramid Network (FPN; Lin et al. (2017a)) to enhance sensitivity to small objects, allowing the model to recognize features at multiple scales and improve detection accuracy across varying target sizes. The model was trained and validated on the MIDOG++ dataset,Aubreville et al. (2023a) a high-quality, multi-domain collection of images spanning various institutions, scanners, and tissue types (both human and animal samples). This diversity strengthens the foundation for training and validating the model, allowing it to learn robust, generalizable features while reducing bias and improving resilience to domain shifts commonly encountered in clinical settings.

Our MF detector demonstrated strong overall performance, achieving an F1-score of 76.36% and an accuracy of 78.13% across all classes, see Table S1. Detection rates varied across tumor types, with true positive rates of 79.95% and 79.35% for Canine Cutaneous Mast Cell Tumor and Human Breast Cancer, respectively. However, higher false negative rates were observed in Human Neuroendocrine Tumor (36.94%) and Canine Lung Cancer (35.52%). These variations underscore the influence of tumor type on detection effectiveness. Despite these challenges, the model achieves a strong balance between precision and recall (high F1-score), ensuring reliable differentiation between positive and negative cases. With further refinement, this model has significant potential for clinical applications, particularly in cancer screening and monitoring, where precise mitotic figure detection is critical for accurate diagnosis and treatment planning.

### 3.2 AES: User-Focussed XAI Query Engine

Our approach focuses on explaining the confidence (“belief score”) produced by an object detection model. This score represents the model’s confidence that an image is a positive case, distinguishing it from methods that focus on the binary positive-negative decision itself, which are commonly emphasized in prototype-based methods Snell et al. (2017). When the detector’s belief/confidence score for an image exceeds the decision threshold value, the image is classified as positive. By analyzing the learned regression function of the confidence scores, we can examine the detection boundary between mitotic and non-mitotic figures and look at actual prototypical examples that inform decision, similarly to how human subject matter experts make judgements when classifying images. Pathologists can gain insights into how a model’s confidence varies when analyzing visually similar images, such as mitotic figures and their “look-alike” non-mitotic counterparts. These insights are crucial for trust and adoption in clinical workflows, where understanding a model’s decision-making process is just as important as the final classification outcome.

In our framework, a pre-trained black-box MF detector processes test histopathological images to generate bounding boxes with associated confidence scores. These scores represent the model’s confidence that a given box contains a mitotic figure. To classify these boxes as mitotic (positive classification) or non-mitotic (negative classification), a confidence threshold is set, optimized to balance false positive and false negative rates. However, the complexity of black-box DL models obscures why certain bounding boxes receive high confidence scores, while others do not. This challenge is compounded at the decision threshold, where the differentiation between mitotic and nearby non-mitotic figures remains unclear, limiting user trust in the binary outcomes.

To address these challenges, our approach provides prototype-based explanations, an approach that is widely regarded as desirable in digital pathology for its ability to combine interpretability with performance. Inspired by foundational work in this area, such as that by Bien and Tibshirani (2011), and modern advancements like ProtoAttend Arık and Pfister (2020), we focus on identifying a concise, predictively relevant set of actual clinical examples (not synthetically generated) to enhance user understanding. In our query engine, bounding boxes predicted as being positive or negative with sufficiently significant confidence scores by the black-box MF detector are used as input. For each input bounding box, the output is a sparse set of active prototypical examples, both mitotic (positive) and nearby non-mitotic (negative) figures, with known class labels. These prototypes represent elements that influenced the decision-making of the black-box model, as determined by a learned regression model. The prototypical examples provide interpretable insights into the model’s decision process, especially at the critical decision boundary (i.e., at the detection threshold value), while minimizing the number of relevant prototypes to improve clarity for end users.

Unlike traditional prototypical methods, which attempt to explain the final binary classification outcomes, our approach emphasizes the confidence scores that inform these decisions. This regression-based strategy is particularly suitable for nuanced medical imaging tasks, such as mitosis detection, where pathologists need to understand how the model’s confidence evolves for similar-looking images. By prioritizing fewer but more informative prototypes and tailoring explanations to end-user needs, our approach ensures that the explainability framework aligns with clinician’s domain knowledge, supporting informed and confident decision-making. To validate this methodology, we trained a black-box DL model and conducted numerical experiments, demonstrating the query engine’s effectiveness in offering meaningful, interpretable explanations of the black-box detector’s outputs, particularly for challenging edge cases.

To demonstrate AES’s interpretability, we present three test scenarios, Figure 3. In these scenarios, the detection threshold is set to *τ*_0_ = 0.969 and the sets of positive and negative decision figures correspond to taking PDF = PDF_*ϵ*_(ℐ) and NDF = NDF_*ϵ*_(ℐ), respectively, for *τ*_*𝓁*_ = 0.85, *ϵ* = 0.01, and 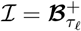.^11^

**Figure 3:**
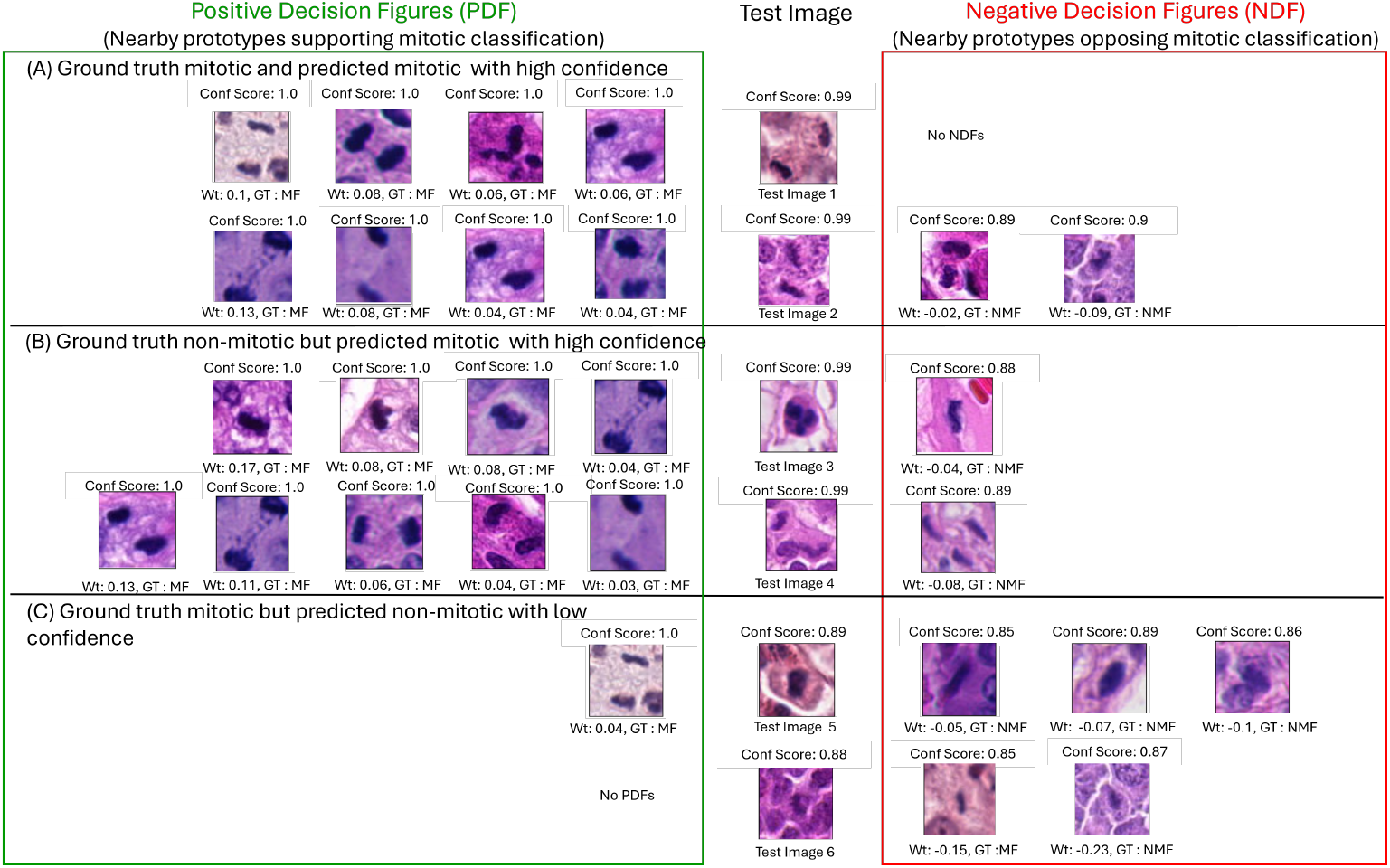
AES-based explanability for mitotic figure detection demonstrated across three test scenarios. For each predicted bounding box, the AES generates nearby PDF and NDF prototypes. These prototypes provide visual insights, allowing users to interpret training or curated examples that influence the MF detector’s decision-making process. For further details, see Section 3.2.

a. **Ground truth mitotic and predicted mitotic with high confidence:** In the first two test cases, the MF detector correctly classifies mitotic figures *x* with confidence scores above the threshold (a confidence score of *β*(*x*) = 0.99 *> τ*_0_). For these cases, we present the active prototypes AES(*x*) identified by a mean relevance score, Mean-RS, chosen to explain approximately 90% of the prediction (see Equations (21)–(23) in the Methods Section). The prototypes on the right correspond to the positive decision figures (PDFs) in AES(*x*) ∩ PDF, while those on the left correspond to the negative decision figures (NDFs) in AES(*x*) ∩ NDF). In the first image, no NDFs are found to influence the mitotic classification. However, in the second image, both PDFs and NDFs contribute to the mitotic classification.
b. **Ground truth non-mitotic but predicted mitotic with high confidence:** In the third and fourth test cases, the MF detector assigns a high confidence score of 0.99 to non-mitotic figures, classifying them incorrectly as mitotic because the score exceeds the threshold of 0.969. AES explains this misclassification by showing nearby mitotic figures (PDFs) that may have misled the model, alongside proximate non-mitotic figures (NDFs).
c. **Ground truth mitotic but predicted non-mitotic with low confidence:** In the fifth and sixth test cases, the MF detector assigns a low confidence score of 0.89 to mitotic figures, classifying them incorrectly as non-mitotic because the score falls below the threshold of 0.969. AES explains this error by identifying nearby non-mitotic figures (NDFs with high weights) that may have influenced the model, while also presenting proximate mitotic figures (PDFs) when available.

These cases highlight AES’s role in enhancing transparency by providing real-world contextual examples, enabling pathologists to effectively audit and interpret model decisions and identify sources of misclassification.

Building on this foundation, our workflow includes user-centric enhancements that improve adaptability and usability for end-users. A key feature is the ability for users to define custom decision boundaries tailored to their specific object detection needs by selecting thresholds on detection confidence scores. This flexibility is critical in object detection systems where classification depends on confidence score thresholds. By allowing users to set these boundaries, our algorithm presents prototypical examples that illustrate how individual cases positively or negatively influence the decision-making process for a given test image.

Further, the user can train the AES algorithm with initial examples/prototypes relevant to their specific use case. This functionality is particularly valuable in histopathology, where experts possess domain-specific knowledge about atypical patterns that may confuse automated systems. For instance, histopathologists can select specific positive and negative decision figures for training our explanatory engine, forcing the AES algorithm to derive coefficients that quantify and visualize the influence of *these* figures on the model’s decisions. These features enhance the algorithm’s adaptability to diverse tasks and provide interpretable insights into its decision-making process, fostering greater transparency and trust among users.

## 4 Conclusions

Developing explainable DL solutions for diagnostic tasks, such as mitosis detection in cancer grading and prognosis, presents significant challenges. In this study, we introduce Adaptive Example Selection, an innovative methodology that offers visual insights into the decision-making process of a well-trained, black-box Faster R-CNN model. AES transforms model interpretability by providing prototypical labeled images that resemble newly classified, previously unseen cases, enabling clinicians to easily understand the model’s decisions. These reference images serve as clear exemplars, improving model transparency, fostering trust, and boosting confidence—key factors for clinical adoption. By focusing on confidence scores and examining decision boundaries, AES goes beyond traditional binary classification explanations. It provides a deeper, more nuanced understanding of the AI model’s behavior, empowering clinicians with the knowledge they need to trust and use AI tools in high-stakes environments.

AES not only improves diagnostic accuracy but also encourages direct clinician-AI collaboration, helping uncover potential model biases and promoting more reliable, equitable healthcare outcomes. With its user-centric design, customizable decision thresholds, and expert-selected prototype integration, our framework is a powerful tool for increasing transparency in medical AI. By ensuring that AI models are not only accurate but also interpretable and adaptable, we take a crucial step toward confidently integrating deep learning into real-world clinical practice.

Our XAI framework has the potential to serve as an automatic diagnostic aid for tumor grading and prognosis, enabling timely treatment decisions and supporting the training of novice pathologists. Looking forward, we plan to extend this work to predict entire ROIs in whole-slide images. Additionally, we intend to conduct a rigorous user study with two to three experienced pathologists to evaluate how our explanation-driven approach affects diagnostic accuracy. This study will compare performance with and without our system’s explanations, assessing its effectiveness in real-world clinical workflows. While this study focused on mitosis detection, the principles behind AES are universally adaptable and are applicable to a wide range of diagnostic tasks, including tumor detection, organ classification, and rare disease diagnosis. Our framework’s flexibility ensures it can be tailored to different clinical environments, contributing to the broader adoption of explainable AI in healthcare.

## 5 Methodology

The Methodology section is organized into two subsections. Subsection 5.1 provides a detailed explanation of the black-box Faster R-CNN MF detector, including its nature, training process, and validation. In Subsection 5.2, we shift focus to the AES query engine, which is trained to analyze the classification decisions made by the MF detector. The AES engine offers visual explanations for each new classification, based on labeled examples known by subject matter experts to be prototypically representative of mitotic or non-mitotic figures.

### 5.1 Black-Box Faster R-CNN MF Detector

#### 5.1.1 Dataset

For our experiment we used the publicly available MItosis DOmain Generalization++ (MIDOG++) dataset Aubreville et al. (2023b), which is the largest multi-domain mitotic figure dataset available at the time of this writing. The dataset consists of high resolution ROI images from 503 histological specimens, containing 11,937 mitotic figures from seven human and canine tumor types with diverse morphologies, including breast carcinoma, lung carcinoma, lymphosarcoma, neuroendocrine tumor, cutaneous mast cell tumor, cutaneous melanoma, and subcutaneous soft tissue sarcoma. These images were digitized by one of 5 whole slide scanners in one of four pathology laboratories. For our experiment, the dataset was divided into training (291 images), validation (79 images), and test (111 images) sets in a 6.5:1.5:2 ratio, ensuring a balanced representation of mitotic figures from each tumor type in both the training and validation sets.

#### 5.1.2 Data Preprocessing

The MIDOG++ dataset contains high-resolution images up to 7215×5412 pixels, but the mitotic instances are very small—only about 50×50 pixels each. To reduce computational demands, DL object detection models typically downsample such large images, often at the cost of fine details. This loss of resolution makes detecting small objects like MFs within large backgrounds particularly challenging. For that, 512×512-pixel image patches were generated using a sliding window with a 20% overlap following the procedure adopted by Aubreville et al. (2023b), thereby ensuring that mitotic objects near patch edges are fully captured. This method increases the relative size of MFs within each patch, from 0.7% of the original image’s side length to about 10%, greatly enhancing detection accuracy in DL models.

To enhance data diversity and improve model generalization in MF detection, we applied various data augmentation techniques. These include random horizontal and vertical flips (50% probability each) and ±45-degree rotations (50% probability) to reduce positional biases and introduce angular variations. To account for lighting variability, brightness and contrast adjustments (20% probability) modify brightness by [−0.2, +0.2] and contrast by ±20%, simulating different imaging conditions. Gaussian noise (variance of [10, 50], 30% probability) further improves robustness, while a 20% probability saturation shift of ±20% manages color variability due to staining differences. Each augmented image is converted to tensor format for model compatibility, creating a diverse dataset that enhances detection accuracy. Additionally, we implemented the Random Stain Normalization and Augmentation framework by Shen and Ke (2022) to align stain color distributions and simulate realistic variations. Four style templates, primarily pink and purple hues, were selected based on visual analysis. K-Means clustering further refined template selection by grouping similar images based on color histograms, ensuring effective capture of staining diversity.

#### 5.1.3 Faster R-CNN: Training and Hyperparameter Tuning

Faster R-CNN is a widely used framework for object detection, especially in natural image datasets like MS-COCO Lin et al. (2015). Built on region proposal methods (RPN) Uijlings et al. (2013) and R-CNNs Girshick et al. (2014), it consists of three interconnected modules: (a) a CNN for feature extraction, (b) an RPN that generates bounding boxes around regions of interest, and (c) an R-CNN that analyzes these bounding boxes to classify each enclosed object. Faster R-CNN treats object detection as both a regression and classification problem. During training, it maintains class balance by selecting a mini-batch from a single image, ensuring a 1:1 ratio of positive to negative anchors before computing the loss. For this study, we employed a Faster R-CNN model with a 50-layer Residual Neural Network (ResNet-50) He et al. (2015) as the feature extractor, pre-trained on MS-COCO dataset which differs from the RetinaNet architecture (Lin et al. (2017b)) used in Aubreville et al. (2023b). To improve sensitivity to small objects, we also integrated a Feature Pyramid Network (FPN) Lin et al. (2017a).

The network was trained on image patches containing mitotic objects, excluding patches without MFs to improve computational efficiency and avoid redundant data. Training was conducted with a batch size of 2, a learning rate of 0.001, and 50 epochs, using stochastic gradient descent with a momentum of 0.9. Validation performance was evaluated by mean average precision (mAP) across Intersection over Union (IoU) thresholds from 0.5 to 0.95 in 0.05 increment. The model achieving the highest mAP was selected for MF detection. After convergence, parameter tuning was conducted to optimize class confidence and object area thresholds for filtering predictions, maximizing the overall F1-score.

The object detection algorithm learns a belief function *β*(·), which maps confidence scores *β*(*x*) ∈ [0, 1] to classify bounding boxes *x* ∈ ℬ ^*all*^. In our binary detection case (mitotic figures versus non-mitotic figures), this function represents the confidence that a bounding box *x* contains a mitotic figure:

*β*(*x*) = confidence that the bounding box *x* contains a mitotic figure; i.e., that *i*_*x*_ ≐ MF.

The set of positively predicted bounding boxes at confidence level *τ*_0_ is given by:

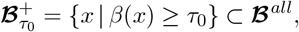

This set contains bounding boxes predicted to contain mitotic figures with at least *τ*_0_ confidence, based on a user-determined threshold *τ*_0_.

#### 5.1.4 Performance Evaluation of the Trained Faster R-CNN

To evaluate the performance of the object detector, we used the following metrics: Precision reflects the proportion of positive classes correctly classified by the model (Eq. 14). Recall indicates the proportion of positive classes correctly identified out of the total positive classes (Eq. 15). F1 score evaluates the balance between precision and recall (Eq. 16). The F1 score shows a strong performance in recognizing positive cases while minimizing false positives and false negatives. This makes it a suitable metric when recall and precision must be optimized simultaneously.

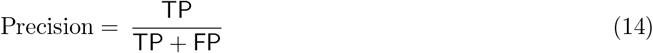

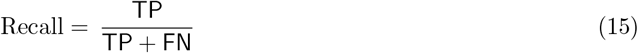

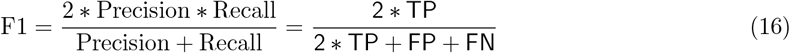

All experiments were conducted on a high-performance Linux server equipped with an NVIDIA RTX A5000 GPU featuring 24 GB of memory, complemented by 64 AMD EPYC 75F3 32-core processors operating at 3.8 GHz and 256 GB of RAM. The software environment was configured with CUDA 11.8 and cuDNN version 9.1, utilizing Python 3.8.10 as the programming language for the implementation. This setup provided an optimal platform for executing computational tasks efficiently. As described in Section 2.1 and detailed in Table 1, a threshold value of *τ*_0_ = 0.969 was determined to provide optimal performance across all tumor types within the dataset.

### 5.2 AES Query Engine

This section assumes familiarity with the notation discussed in Sections 2.2.1 and 5.1.3.

The main objective of the AES framework is to interpret and explain the confidence scores function *β*(·) of predicted bounding boxes from the trained MF detector. As a prototype-based XAI technique, AES aims to closely approximate these confidence scores while providing visual insights by highlighting the influence of both mitotic and non-mitotic figures in close proximity, referred to as a set of active prototypes. In this study, active prototypes are drawn from the training data, but can potentially be replaced with specific examples provided by subject matter expert.

The framework works in two distinct steps. In the first step, a test histopathology image patch is passed through the MF detector’s inference engine, producing a collection of candidate bounding boxes 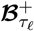. The threshold *τ*_*𝓁*_ can be selected (as discussed below) to filter out boxes unlikely to contain mitotic figures or based on other user-defined criteria. For each bounding box 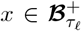, the value *β*(*x*) ∈ [0.1] serves as a measure of confidence that bounding box *x* envelopes an image *i*_*x*_ that contains a mitotic figure. Future work will explore conformalization of these confidence measures to ensure that they are well calibrated, e.g., to empirical false positive rates Barber et al. (2023).

In the next step, the XAI query engine works as follows. For each bounding box 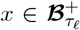, the AES query engine generates a small set of labeled clinical images, denoted as AES(*x*) which serve as active prototypes^12^. These active prototypes provide a concise and informative approximation 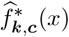 of the transformed confidence score (see Eq. (4)) *f* (*x*) = SBC(*β*(*x*); *τ*_0_), given by

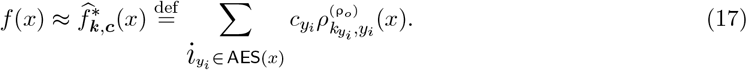

This set AES(*x*) offers visual insights into how the MF detector assigns confidence scores to bounding boxes *x*. It is important to note the distinction between the concisely informative approximation 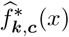 and the more general approximation 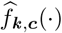 of Eq. (5).

The AES approximation process begins with pre-processing and sorting the data, the predicted confidence scores, and the potential set of active prototypes, which is discussed in Sections 5.2.1 and 5.2.2. Following this, Section 5.2.3 describes the RBF basis functions used to approximate the transformed confidence scores. In Section 5.2.4, we provide a detailed explanation of the loss terms in the optimization algorithm and how they contribute to achieving our explainability goals. Finally, Section 5.2.5 outlines the metrics used to evaluate the effectiveness of the AES methodology.

#### 5.2.1 Pre-processing and Sorting the Data Using Learned Confidence Scores

The initial pre-processing step involves removal of predicted bounding boxes with lower areal sizes and overlapping predictions. Let ℬ ^*all*^ be the set of all predicted bounding boxes determined from the MF detector that meet specific admissibility criteria. These criteria include: an areal size of at least 2400 pixels (motivated by the provided squared approximated bounding boxes of equal size (50 × 50 pixels) in the MIDOG++ dataset) and filtering of the data via an application of non-maximum suppression (NMS) to eliminate redundant overlapping detections.

Next, the set ℬ ^*all*^ is further sorted into subsets based on their confidence scores *β*(*x*) to be utilized for *Decision Boundary Analysis* (DBA) in the following section. Specifically, we define two sets:

- The set of at least *τ*_0_-level positively-predicted bounding boxes by

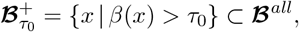

for a user-determined confidence threshold *τ*_0_.
- The set of general bounding boxes

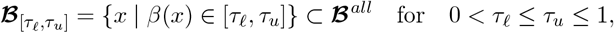

which aims to incorporate user-defined thresholds and obtain a nuanced understanding of the model’s decision boundary by considering both positive and negative examples in close proximity to the threshold value *τ*_0_. In particular, if we take

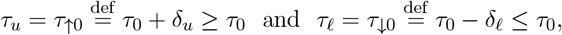

for small *δ*_*u*_ ≥ 0 and *δ*_*𝓁*_ ≥ 0, then we can interpret ℬ_[*τ* ↓0,*τ*↑0]_ as the set of positive and negative decision “look-alikes” relative to the positive-decision threshold *τ*_0_. Note that taking *δ*_*u*_ = 0 gives the set, ℬ_[*τ*↓0,*τ*0]_, of “almost-positive decision” bounding boxes *x*.

Instead of examining only the predicted bounding boxes 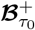, we incorporate the almost-positive boxes and focus on the enlarged set

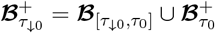

where the bounding boxes *x* in 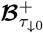 contain both the predicted mitotic figures *i*_*x*_ at level 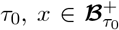, and the nearby (at decision level *τ*_*𝓁*_ = *τ*_0_ − *δ*_*𝓁*_) almost-positive negative figures *i*_*x*_′ for *x*^′^ ∈ ℬ_[*τ*↓0,*τ*0]_. By focusing in detail on the elements in 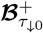 we can find prototypical cell images that inform clinicians about the granular details that the trained decision algorithm uses to distinguish between mitotic versus non-mitotic images. This enables a clinician to establish a personal degree of trust in the algorithm as well as allowing them to provide experience-based feedback to the algorithm that can be used to improve its trust and performance.

Based on cross-validation on the MF detector model (optimizing on F1 metric for detecting mitotic versus non-mitotic cells), we established a decision threshold of *τ*_0_ = 0.969 ∼ 96.9% on the confidence scores; if a bounding box’s confidence score exceeds this threshold the prediction is labeled a positive (i.e., that the box 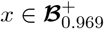 contains a mitotic figure at belief level 96.9%) and otherwise is labeled negative.

We also take *τ*_↓0_ = 0.850 ∼ 85.0% which corresponds to setting *δ*_*𝓁*_ = 0.119 ∼ 11.9%.^13^ This yields

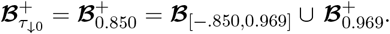

Note that 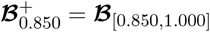. The distribution of the confidence scores *β*(*x*) for the members of the set 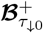 can be highly skewed and bunched together, as shown in the lower left of Figure 2 for 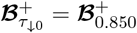 determined from the MIDOG++ data with the confidence score level set at *τ*_0_ = 0.969 ∼ 96.9. This makes it difficult to examine the structure of cell images continued in the bounding boxes with scores around the level of the baseline confidence score. As introduced in Section 2.2.2 and described in 5.2.2, we apply variable-stabilizing power transformations on the confidence score function to expand the behavior of the MF detector’s classification particularly around the decision threshold *τ*_0_, which we refer to as DBA.

#### 5.2.2 Decision Boundary Analysis: “Zooming in” and Expanding the Belief Function

The inclusion of DBA enhances the AES algorithm by magnifying the decision boundary determined by the confidence scores *β*(*x*) around the threshold *τ*_0_. It achieves this by stabilizing variance and employing appropriate sign constraints, allowing the AES algorithm to effectively contrast new test images with nearby mitotic and non-mitotic bounding boxes in 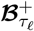.

The rationale behind DBA is to query the decisions by zooming into the vicinity of the threshold boundary *τ*_0_, where the algorithm is least likely to make a decision with confidence, to identify image types that tend to confuse the detector. Since 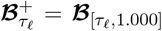, this approach also allows for analyzing highly confident mitotic classifications (i.e., images *i*_*x*_ for which *β*(*x*) ≈ 1) to determine to what degree this confidence is well-founded.

To facilitate this “expanded view” of decision making near the decision threshold value *τ*_0_, we transform the confidence scores *β*(*x*) in two steps: First, they are transformed using a variance-stabilizing Box-Cox (BC) transformation; Second, they are then centered at the transformed value of the detection threshold *τ*_0_. This results in centered BC-transformed confidence scores *f* (*x*).

The BC transformation of a nonnegative real variable *y* is given by:

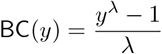

We define a shifted BC transformations as:

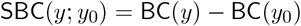

for some fixed, nomimal value *y*_0_.

We transform the MF detector’s confidence scores *β*(*x*) ∈ [0, 1] via

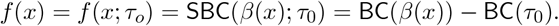

The optimal parameter for *λ* is estimated through maximum likelihood using this [package](https://scikit-learn.org/1.5/modules/generated/sklearn.preprocessing.power_transform.html). Notably, *f* (*x*) = 0 at the decision threshold value *β*(*x*_0_) = *τ*_0_. In that case, *x* represents a *positive decision figure* for a given threshold value *τ*_0_ if *f* (*x*) *>* 0 and represents a *negative decision figure* otherwise.

#### 5.2.3 Approximating the Expanded Belief Function via Truncated Radial Basis Functions

The learned function 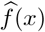 of Eq. (5) approximates the shifted BC transformation *f* (*x*) = SBC(*β*(*x*)) of Eq. (4). To achieve this, we first extract latent representations *r*_*x*_ for bounding boxes *x* from the penultimate layer of the MF detector for the set ℳ_train_ during training and from the set ℳ_test_ during testing. Then we normalize them by projecting into directional vectors given by the transformation 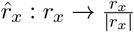.

For any two bounding boxes *x* and *y* in ℬ ^*all*^, we define a divergence measure:

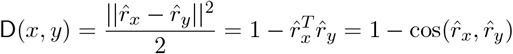

where 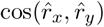 represents the cosine similarity between the normalized representations 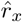 and 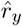. For a fixed value of *y*, we then have the RBF *centered at* 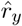 (or, for short, *centered at y*) defined by^14^

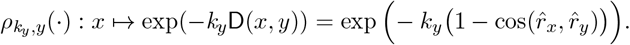

Note that other divergence or distance measures (such as Euclidean norms) result in a good approximation of *f* (*x*) or facilitates interesting exploration of the geometry of the embedding space can also be applied here.

In our situation |D(*x, y*)| ≤ 1, therefore the RBFs 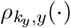 have finite (and hence compact) support as a function of the penultimate-layer representation vectors *r*_*x*_ and *r*_*y*_. Further, in general *k*_*y*_ ≠ *k*_*y*_′ for *y* ≠ *y*^′^ so that, in general, the RBF is *not symmetric* with respect to the interchange of *x* and *y*,^15^

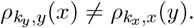

Note that only for the special case when *k*_*y*_ is independent of *y* (i.e., when *k*_*y*_ = *k* = constant) do we expect the RBF to be generally symmetric.^16^

To further enforce concentrated localization of the finite-support RBF (7) about its center point *y*, the values of the RBF basis functions are trimmed (truncated to zero) according to the value of a trim parameter ρ_*o*_ in the following way:

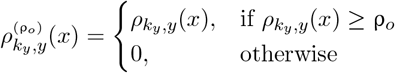

for 0 ≤ ρ_*o*_ ≤ 1. The relationship between the concentration parameters *k*_*y*_ and trim parameter *ρ*_0_ is explored in the Supplementary section S4.

#### 5.2.4 Goals of the AES Training Optimization of Section 2.2.4

Having established the pre-processing steps and the RBF basis functions, we now summarize the key objectives of our optimization framework. To ensure good behavior of the AES query engine, our optimization is designed to:

1. Learn a small yet informative set of candidate active prototypes 𝒟 whose elements *y*_*i*_ ∈ 𝒟 comprise the RBF centers *y*_*i*_ used in our approximation 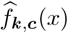 of the expanded belief function *f* (*x*) and from which active prototypes (the elements of AES(*x*)) will be selected to provide visual insight into the classification of image *i*_*x*_.
2. Ensure that the RBF centers *y*_*i*_ ∈ 𝒟 are well separated and spread-out to ensure that the small collection of centers comprising the set 𝒟 have wide and informative coverage over every possible image that could be encountered in the future.
3. Produce large values of the concentrations 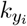 to further enhance localization of the RBFs, thereby forcing the RBFs to be placed on well-positioned centers *y*_*i*_ ∈ 𝒟;
4. Learn values of *c*_*i*_ that ensure a good approximation 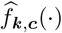 to *f* (·) while aligning the signs of the values of *f* (*x*) with their respective approximation values 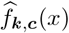 in order to further promote consistency between the decisions obtained from the values *f* (*x*) and the decisions obtained from the values 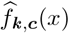.

These goals are by no means mutually exclusive or orthogonal to each other. Below, we discuss these goals and how they are reflected in the choice of the loss functions of Equations 9 and 10 and the rationale behind their alternating optimization.

Starting with a initial subset of example prototypes of interest 𝒟_0_, we want to obtain a reduced set of active prototypes 𝒟 which after optimization is a refinement of the initial set 𝒟 _0_. The initial set 𝒟 _0_ might be selected with the help of pathologists’ preferred prototypes relevant to a problem domain. The examples in 𝒟 (or 𝒟 _0_) are separated into two groups :

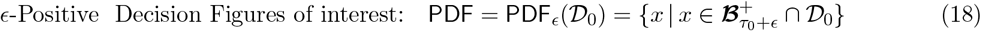

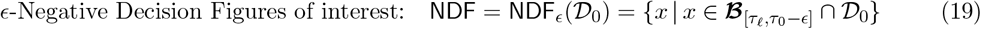

PDF_*ϵ*_ active prototypes represent cases where the MF detector predicts mitotic, while NDF_*ϵ*_ active prototypes indicate cases classified as non-mitotic. This approach aims to guide the learning process of the AES model towards more relevant relationships in the data by incorporating prior knowledge, enhancing interpretability, and ensuring that the parity of the signs is meaningful. The selection of PDF_*ϵ*_ and NDF_*ϵ*_ is flexible; for e.g., a subject matter expert might want to focus on true positives or true negatives rather than all figures. Further they can also choose specific prototypes based on their domain knowledge that assist in interpreting the relationship with the detector’s decision-making process. This flexibility allows SMEs to tailor the model’s focus to their specific analytical needs and research objectives.

Formally, we want the set of active prototypes 𝒟 ⊂ *X* to satisfy the following conditions,

1. **1. Sign-Alignment**: The approximation of the transformed belief score *f* (*x*) is given by Eq. (5), where *c*_*i*_ ≥ 0 for *y*_*i*_ ∈ PDF_*ϵ*_ and *c*_*i*_ ≤ 0 for *y*_*i*_ ∈ NDF_*ϵ*_.
2. **2. Spread-Out Sparsity**: The cardinality of the set of global prototypical examples 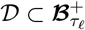 is much lower than the cardinality of 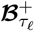, ensuring succinctness (sparsity) of representation. Further encouragement of small values of the condition number *κ*(𝒟) results in spreading-out of the prototypes *y*_*i*_ ∈ 𝒟 which assists in the avoidance of multicollinearity of the prototypes.
3. **3. Localization** (Fewer Relevant Examples per Prediction): The value of 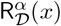 (see Eq. S9) is small for all *x*, ensuring that fewer relevant examples AES(*x*) ⊂ 𝒟 are selected for explaining the predicted classification of a datum *x*.

These conditions serve to enhance interpretability by aligning with prior PDF_*ϵ*_ and NDF_*ϵ*_ knowledge, reduce redundant examples, and provide sparse, robust interpretations for end-users. The sign-alignment condition minimizes cancellation effects (by imposing meaningful parity of signs), while sparsity and localization ensure an efficient and focused prototype selection, both globally as well as for individual predictions.

The AES model performs functional approximation (employing two neural networks) of the confidence scores with the RBF feature vectors. It utilizes an alternate coordinate descent trained with the loss functions given by Equations 9 and 10.

- The first loss function is given by:

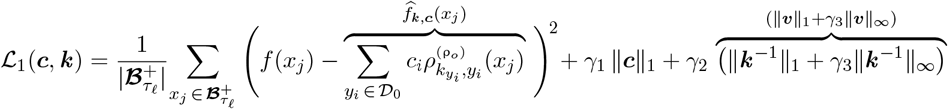 This loss function consists of four components:
  - A regression quadratic-loss term

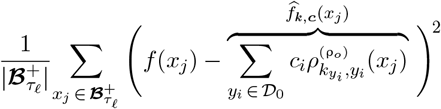

giving a sample-averaged squared-error between the transformed confidence scores *f* (·) and an approximation 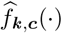 formed from a weighted linear combination of the feature vectors 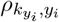. The sets 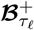 and 𝒟_0_ are constructed from training data provided by the previously trained Faster R-CNN decision engine.
  - The term ∥***c***∥_1_ provides a 1-norm regularization on the coefficients *c*_*i*_ associated with the RBF centers *y*_*i*_ ∈ 𝒟_0_, thereby promoting sparsity of 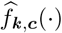.
  - The term ∥***k*** ^−1^∥_1_ = ∥***v***∥_1_ enforces a 1-norm regularization on the variability parameters 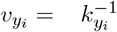 associated with the RBF centers *y*_*i*_ ∈ 𝒟_0_, thereby enforcing many small scale values 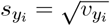 (equivalently, large concentrations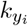).
  - The term ∥***k*** ^−1^∥_∞_ = ∥***v***∥_∞_ = ∥***s***^2^∥_∞_ ensures that the the larger scale parameters 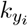 (the ones not forced to be small as a consequence of the one-norm regularization) are bounded, which helps localize the influence of the prototypes.
- The second loss function is defined as:

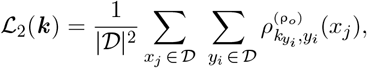

and it operates solely on the pruned or active prototypes 𝒟. This loss function ensures that the RBFs associated with the prototypes in 𝒟 remain distinct (i.e., have little overlap) by adjusting their concentration (equivalently, scale) parameters.

The loss functions are alternately trained for 10 epochs each, with early stopping based on either the ℒ_1_ validation loss or when the *R*^2^ score drops below 0.995 times the maximum *R*^2^ score observed on the validation data during training. The loss functions have been trained using PyTorch’s DL frameworks. The training and test splits follow the same partitioning used in the Faster R-CNN model, with 358 training images and 111 test images. The metrics reported in Table 2 are computed on these 111 test images, considering all admissible bounding boxes with a confidence of at least 85%.

#### 5.2.5 Metrics for Prototype Evaluation

The learned dictionary 𝒟 provides a globally sparse representation of the training dataset 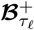 which (if the training data set is sufficiently representative and our training methodology sufficiently robust) should also be representative of any properly sampled test dataset 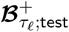. Consider the trained model approximating the normalized belief scores:

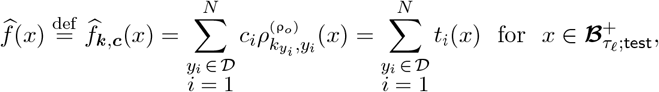

where #Act= *N* = |𝒟| is the cardinality of the set of learned active prototypes 𝒟. The *R*^2^-score measures how well the predictions from 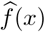, derived from the learned dictionary 𝒟, approximate the normalized scores *f* (*x*). For a fixed representation vector *x* selected from 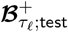, and the terms, indexed as *t*_*i*_(*x*), *i* ∈ ***I*** = {1, · · ·, *N* = |𝒟 |}, are defined by

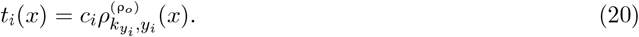

For a selected fixed value of *x*, we uniquely sort the absolute values |*t*_*i*_(*x*)| in decreasing order of magnitude and define a re-indexing *j* ∈ ***J*** = {1, · · ·, *N* = |𝒟|} via a permutation: 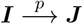 according to

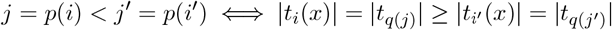

where *q*(·) is the inverse permutation to *p*(·), *q*(*j*) = *q*(*p*(*i*)) = *i* ∈ ***I*** and *p*(*i*) = *p*(*q*(*j*)) = *j* ∈ ***J*** .

Set *t*_0_(*x*) ≡ 0 and define the partial-sum approximation,

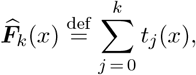

to be the sum of contributions of the *k* largest absolute value terms predicting 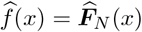. We define a *α*-parameterized *k*-loss function for *α* ∈ [0, 1] as follows:

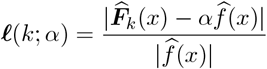

We define the *partial-sum relevance* at level *α* for a particular input 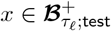 as,

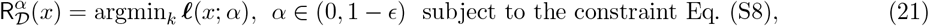

which provides an integer value 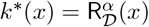 that gives the minimum number of active prototypes (the number of terms in the optimal partial-sum approximation 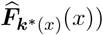 needed to estimate 100*α* percent of the total-sum approximation 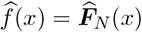. Once *k*^*^(*x*) is at hand we can form a set of locally active prototypical examples that inform the classification decision for the vector *x*:

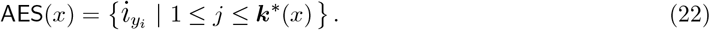

Note that this is equivalent to Eq. (13) when one sets 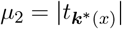. Further, it is evident that 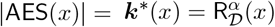.

We then define the *sample mean statistic* Mean-RS *at percent level* 100*α*% by

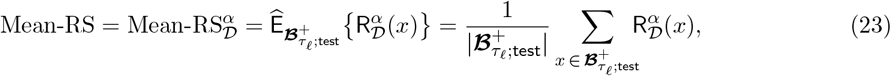

where 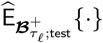 represents a sample average over the test data set 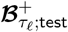. Similarly, we define the *sample median statistic* Med-RS *at percent level* 100*α*% by

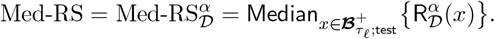

We want these measures to be small, so that only a few number of informative prototype examples appear at the time of inference on a new datum *x*. Notably, the Med-RS score conceptually aligns with the median number of prototypes used in decision-making, as discussed in Arık and Pfister (2020).

### Data

Faster R-CNN pretrained model ℱ

‘look-alike’ threshold *τ*_*𝓁*_ = 0.85;

detection threshold *τ*_0_ = 0.969;

Positive Decision Figure (PDF) and Negative Decision Figure (NDF) initial prototypes.

### Hyperparameters

*γ*_1_, *γ*_2_ and *γ*_3_.

### Result

Trained XAI model - AES

### Pre-processing steps

- Collect latent space representations from the penultimate layer (maybe replaced by other layers) of ℱ for predicted admissible bounding boxes of the training data : 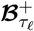 and PDF and NDF figures: *X* = {*z* | *z* ∈ PDF ∪ NDF}.
- Calculate the transformed confidence scores on 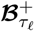 by first applying Box-Cox and then centering the distribution using *τ*_0_. These transformed scores will serve as the target variable, denoted as *y*_true_.
- For each *z ∈* PDF *∪* NDF, create vectors *d*_*z*_ : *x 1→ d*(*x, z*), where *d*(*x, y*) is the cosine distance between *z* and 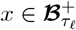.

### Learnable parameters

Concentration parameters *k*_*z*_ and regression coefficients *c*_*z*_ such that 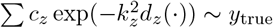

#### Algorithm 1

AES algorithm

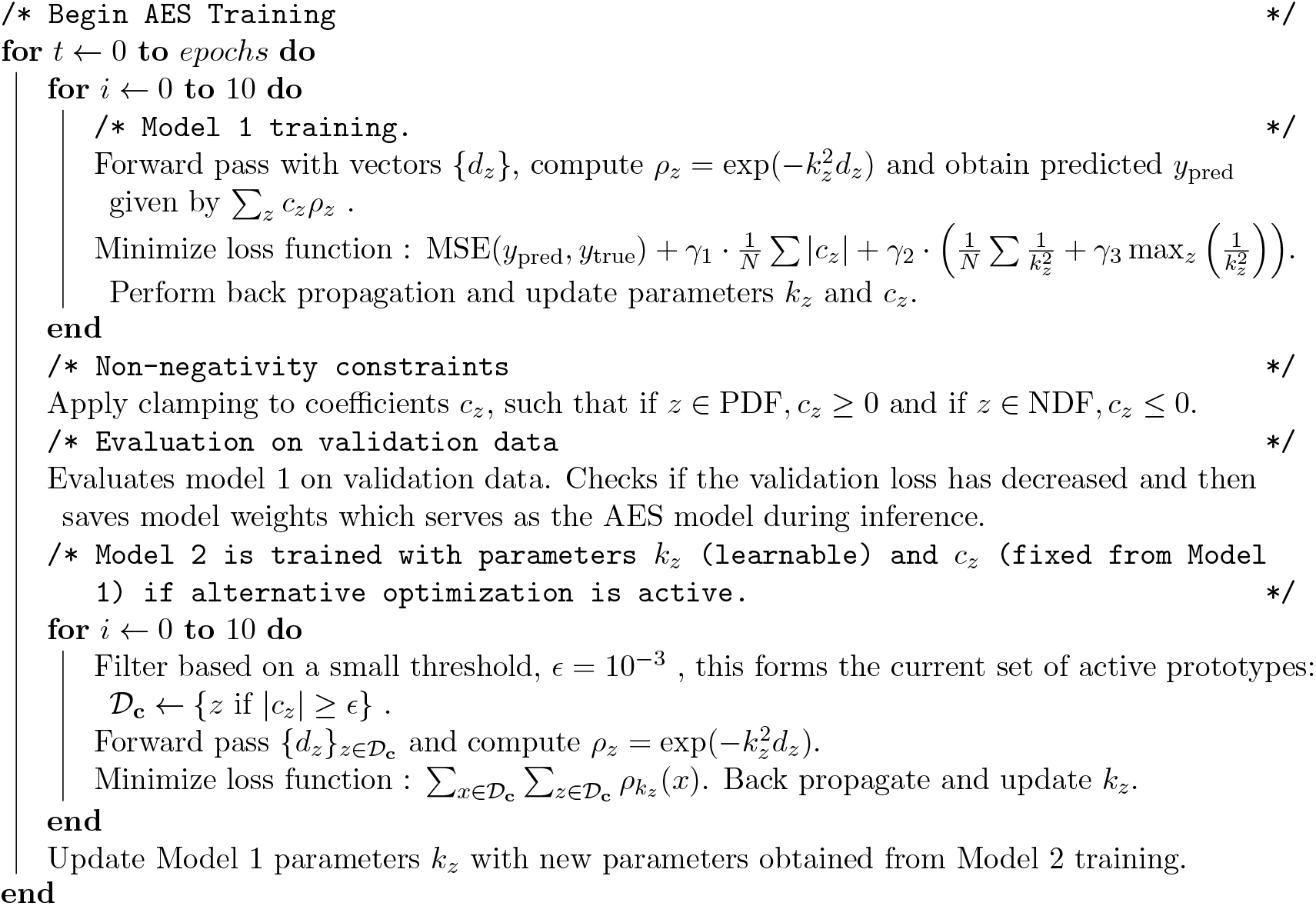

## 6 Data availability

The code that support the findings of this study is available upon reasonable request.

## 7 Author Contributions

**MB:** Investigation (lead); Methodology (lead); Conceptualization (equal); Writing - Original Draft (equal), Review, and Editing.

**KKD:** Writing - Original Draft (equal), Review, and Editing; Methodology (supporting); Investigation (supporting); Supervision (supporting).

**IM:** Methodology (supporting); Writing - Original Draft (supporting), Review, and Editing.

**BB:** Writing - Review and Editing.

**NS:** Supervision (lead); Writing - Original Draft (equal), Review, and Editing; Conceptualization (equal); Methodology (supporting)

## 8 Acknowledgments

We would like to thank Laura Oiye for her invaluable assistance in proofreading and providing feedback on the manuscript.

## Footnotes

1 The default is to just set ℐ= ℬ ^*all*^.

2 See the discussion in the Methods Section.

3 And, for convenience, define ***v***^−1^, ***s***^2^ via component-wise operations.

4 As discussed below, these two optimizations over ***k*** are not independent. To obtain a good estimate of the model parameters, early stopping of the ℒ _2_ minimization is performed.

5 i.e., for each *i* the optimization attempts to learn an 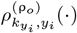that is sharply peaked about *y*_*i*_. Spread and isolation of the centers *y*_*i*_ is also affected by the user specified choices of the support truncation parameter ρ_0_ and the elements of the set 𝒟_0_ used to initialize the optimization algorithm (9)–(10).

6 For sharply peaked RBFs, setting *x* = *y*_*i*_ in Eq. (5) gives 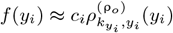. Thus, consistency requires that sign(*c*_*i*_) = sign(*f* (*y*_*i*_)). For the set of all positively identified mitotic images, this corresponds to a nonnegative factorization of the system Eq. (17), and similarly—up to a global sign change—for negatively classified images.

7 The condition number *κ*(𝒟) ∈ [1, ∞) is defined as the ratio of the largest to the smallest singular values of the | 𝒟| × | 𝒟| matrix *AA*^*T*^, *A* = **RBF**(𝒟), where the elements of the 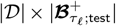 generically full row-rank matrix *A* are computed using centers *y*_*i*_ ∈ 𝒟 and concentration parameters 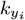 learned from training data ℳ_train_. The model is then evaluated on test Data 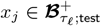. A lower values of *κ*(D) indicates better conditioning of the matrix.

8 The mean statistics presented here are derived from 10 distinct runs. For detailed information on standard errors and AES hyperparameters, refer to the Supplementary Table S2.

9 The *γ*_1_ enforces sparsity of ***c***, while *γ*_2_ minimizes the variability vector ***v*** (equivalently, of the scale vector ***s***) ensuring that the RBFs maintain tight support. The addition of the *γ*_3_ regularization term helps to ensure that the RBFs don’t become Kronecker delta functions by preventing the supports from becoming excessively tight.

10 RBFs are often used for approximation in high dimensional space Lee et al. (1999) and when the scale is constant, quadratic loss-function approximations are closely related to learning in Reproducing Kernel Hilbert Spaces (RKHS).

11 See Eq.s (18) and (19).

12 We refer to the images contained in this set as the *active prototypes* for the bounding box *x*.

13 Lowering the cut-off or detection threshold boosts recall while reducing precision and, as noted above, this allows us to capture more false negatives near true positive figures. The threshold, denoted as *τ*_*𝓁*_, was selected to achieve a recall of around 85% and F1 score of about 70% on the test data. This enables the AES model to explain the misclassification of mitotic figures close enough to the threshold.

14 Note that 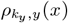 is nonnegative and that its maximumm is achieved: 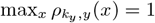.

15 Note, however, that 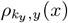 *as a function of x only* is symmetric about the center point *y*. The fact that 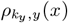 is not symmetric with respect to interchange of *x* and *y* means it is not a kernel function, and thus the framework and tools of reproducing kernel Hilbert space (RKHS) theory are not available for use.

16 If the scale parameters are independent of *y*, so that *k*_*y*_ = *k* for all *y*, then the RBF *ρ*_*k*,*y*_(*x*) is symmetric with respect to an interchange of *x* and *y*. In this case the *kernel matrix* 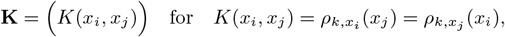 is symmetric and positive semi-definite.^17^ *K*(*x, y*) is therefore a kernel function and induces a Reproducing Kernel Hilbert space (RKHS) as a consequence of the Moore–Aronszajn theorem. Because we work with *y*-dependent scale parameters, in our methodoloogy the matrix **K** is not generally symmetric and we cannot invoke the machinery of RKHS theory.

