## Supplementary Information for "Adaptive Example Selection: Prototype-Based Explainability for Interpretable Mitosis Detection"

March 5, 2025

### S1 Confusion matrix for the MF detector

The confusion matrix for the trained Faster R-CNN model is presented in Table S1.

| **Tumor Type** | **FP** | **FN** | **TP** |
|---|---|---|---|
| Human Breast Cancer | 98 | 70 | 269 |
| Canine Lung Cancer | 55 | 76 | 138 |
| Canine Lymphosarcoma | 144 | 231 | 600 |
| Canine Cutaneous Mast Cell Tumor | 36 | 97 | 387 |
| Human Neuroendocrine Tumor | 66 | 41 | 70 |
| Canine Soft Tissue Sarcoma | 65 | 57 | 190 |
| Human Melanoma | 47 | 47 | 171 |

Table S1: Confusion Matrix for the MF detector

### S2 Simulations from the AES framework

The following simulations, each performed for 10 seeds, were conducted for $\lambda = 23.23$, which was determined using the Box-Cox (BC) Transformation, with a patience value of 10 for the AES model convergence.

| scale | trim | $\gamma_1$ | $\gamma_2$ | $\gamma_3$ | $R^2$ | Mean-RS$^{\alpha=0.9}$ | Med-RS$^{\alpha=0.9}$ | #Act | $\log_{10}(\kappa(\mathcal{D}))$ |
|---|---|---|---|---|---|---|---|---|---|
| constant | 0 | 0 | 0 | 0 | $0.97 \pm 0.00$ | $606.95 \pm 23.91$ | $146.10 \pm 4.89$ | $1647.50 \pm 63.87$ | $5.70 \pm 0.08$ |
| constant | 0 | 1 | 2 | 1 | $0.95 \pm 0.00$ | $253.93 \pm 4.82$ | $238.90 \pm 5.69$ | $1576.20 \pm 11.38$ | $4.93 \pm 0.03$ |
| variable | 0 | 1 | 2 | 1 | $0.96 \pm 0.01$ | $268.85 \pm 153.52$ | $245.90 \pm 99.60$ | $620.90 \pm 313.93$ | $4.46 \pm 0.35$ |
| constant | 0.1 | 0 | 0 | 0 | $0.96 \pm 0.00$ | $621.50 \pm 13.91$ | $791.00 \pm 20.21$ | $1619.10 \pm 23.50$ | $4.16 \pm 0.02$ |
| constant | 0.1 | 1 | 2 | 1 | $0.94 \pm 0.00$ | $182.03 \pm 3.59$ | $210.60 \pm 4.47$ | $1844.50 \pm 17.60$ | $4.36 \pm 0.03$ |
| variable | 0.1 | 1 | 0 | 0 | $0.97 \pm 0.00$ | $530.06 \pm 25.61$ | $690.30 \pm 29.05$ | $1403.30 \pm 76.56$ | $4.02 \pm 0.07$ |
| variable | 0.1 | 1 | 2 | 1 | $0.96 \pm 0.00$ | $21.17 \pm 0.35$ | $8.5 \pm 0.48$ | $269.40 \pm 4.65$ | $2.96 \pm 0.04$ |
| $\mathcal{L}_2 \neq 0$ | | | | | | | | | |
| adaptive | 0.1 | 1 | 2 | 1 | $0.96 \pm 0.00$ | $14.81 \pm 0.36$ | $10.00 \pm 0.49$ | $190.20 \pm 3.07$ | $2.87 \pm 0.05$ |

Table S2: Mean and standard errors from AES Simulations with BC Transformed Confidence Scores (extension of Table 2).

We experimented by turning off the alternative optimization and performed optimization with by additively combining the two loss functions,

$$\mathcal{L}(\boldsymbol{c}, \boldsymbol{k}) = \mathcal{L}_1(\boldsymbol{c}, \boldsymbol{k}) + \eta\, \mathcal{L}_2(\boldsymbol{k}).$$

These experiments were all performed for variable radius framework with trim$= 0.1$,$\gamma_1 = 1, \gamma_2 = 2$, $\gamma_3 = 1$ with patience 10 each for 10 seeds, presented in Table S3 below.

### S3 On the determination of the set of locally active prototypes $\mathsf{AES}(\boldsymbol{x})$

The learned dictionary $\mathcal{D}$ provides a globally sparse representation of the training data set $\boldsymbol{\mathcal{B}}^+_{\tau_\ell}$ which (if the training data set is sufficiently representative and our training methodology sufficiently robust)

| $\eta$ | $R^2$ | Mean-RS$^{\alpha=0.9}$ | Med-RS$^{\alpha=0.9}$ | #Act | $\log_{10}(\kappa(\mathcal{D}))$ |
|---|---|---|---|---|---|
| 0.1 | 0.96 ± 0.00 | 21.18 ± 0.49 | 8.00 ± 0.47 | 268.70 ± 3.64 | 2.98 ± 0.07 |
| 0.5 | 0.96 ± 0.00 | 18.66 ± 0.48 | 6.80 ± 0.39 | 244.00 ± 3.86 | 2.97 ± 0.05 |
| 1.0 | 0.96 ± 0.00 | 18.38 ± 0.61 | 7.00 ± 0.58 | 244.1 ± 3.18 | 2.92 ± 0.07 |
| 2.0 | 0.95 ± 0.00 | 17.68 ± 0.85 | 10.40 ± 0.92 | 245.40 ± 3.42 | 2.87 ± 0.06 |

Table S3: Mean and standard errors for simulations with additive losses and alternative optimization turned off.

should also be representative of any properly sampled test data set $\boldsymbol{\mathcal{B}}^+_{\tau_\ell;\mathsf{test}}$. Consider the trained model approximating the normalized belief scores,

$$\widehat{f}(x) \stackrel{\text{def}}{=} \widehat{f}_{\boldsymbol{k},\boldsymbol{c}}(x) = \sum_{\substack{y_i \in \mathcal{D} \\ i=1}}^{N} c_i \rho^{(\rho_o)}_{k_{y_i},y_i}(x) = \sum_{\substack{y_i \in \mathcal{D} \\ i=1}}^{N} t_i(x) \quad \text{for} \quad x \in \boldsymbol{\mathcal{B}}^+_{\tau_\ell;\mathsf{test}}, \tag{S1}$$

where $N = |\mathcal{D}|$ is the cardinality of the set of learned active prototypes $\mathcal{D}$, $x$ is a fixed representation vector selected from $\boldsymbol{\mathcal{B}}^+_{\tau_\ell;\mathsf{test}}$, and the terms, indexed as $t_i(x)$, $i \in \boldsymbol{I} = \{1, \cdots, N = |\mathcal{D}|\}$, are defined by

$$t_i(x) = c_i \rho^{(\rho_o)}_{k_{y_i},y_i}(x). \tag{S2}$$

For a selected fixed value of $x$, we uniquely sort the absolute values $|t_i(x)|$ in decreasing order of magnitude and define a re-indexing $j \in \boldsymbol{J} = \{1, \cdots, N = |\mathcal{D}|\}$ via a permutation $\boldsymbol{I} \xrightarrow{p} \boldsymbol{J}$ according to

$$j = p(i) < j' = p(i') \iff |t_i(x)| = |t_{q(j)}| \geq |t_{i'}(x)| = |t_{q(j')}| \tag{S3}$$

where $q(\cdot)$ is the inverse permutation to $p(\cdot)$, $q(j) = q(p(i)) = i \in \boldsymbol{I}$ and $p(i) = p(q(j)) = j \in \boldsymbol{J}$.

In the remainder of this subsection the parameters indexed $j \in \boldsymbol{J}$, are always taken as conforming to the ordering of the terms by absolute magnitude according to Eq. (S3). Furthermore, the terms are considered to have been renamed (re-indexed) via

$$t_j(x) \leftarrow t_i(x) = t_{q(j)}(x) \quad \text{for} \quad j \in \boldsymbol{J} \quad \text{and} \quad i = q(j) \in \boldsymbol{I}$$

so that henceforth in this subsection,

$$j \leq j' \iff |t_j(x)| \geq |t_{j'}(x)|. \tag{S4}$$

Set $t_0(x) \equiv 0$ and define the partial-sum approximation,

$$\widehat{\boldsymbol{F}}_k(x) \stackrel{\text{def}}{=} \sum_{j=0}^{k} t_j(x),$$

to be the sum of contributions of the $k$ largest absolute value terms predicting $\widehat{f}(x) = \widehat{\boldsymbol{F}}_N(x)$. Because the terms $t_j(x)$ can change sign, the partial-sum approximation $\widehat{\boldsymbol{F}}_k(x)$ generally is not a monotonic function of $k$. We also define the complementary partial sum,

$$\widehat{\boldsymbol{F}}^c_k(x) = \sum_{j=k+1}^{N} t_j(x).$$

Note that

$$\widehat{\boldsymbol{F}}_N(x) = \widehat{\boldsymbol{F}}_k(x) + \widehat{\boldsymbol{F}}^c_k(x) \;\text{ and }\; \widehat{\boldsymbol{F}}_0(x) = \widehat{\boldsymbol{F}}^c_N(x) = 0.$$

We define a $\alpha$-parameterized $k$-loss function for $\alpha \in [0,1]$ as follows:[1,2]

$$\boldsymbol{\ell}(k;\alpha) = \frac{|\widehat{\boldsymbol{F}}_k(x) - \alpha \widehat{f}(x)|}{|\widehat{f}(x)|} = \frac{|\widehat{\boldsymbol{F}}_k(x) - \alpha \widehat{\boldsymbol{F}}_N(x)|}{|\widehat{\boldsymbol{F}}_N(x)|} = \left| \frac{\widehat{\boldsymbol{F}}_k(x)}{\widehat{\boldsymbol{F}}_N(x)} - \alpha \right| \geq 0. \tag{S7}$$

Note that this can be as small as possible with respect to $k$, $\boldsymbol{\ell}(k;\alpha) \approx 0$,[3] only if there exists a value of $k$ for which

$$\widehat{\boldsymbol{F}}_k(x) = \alpha \widehat{\boldsymbol{F}}_N(x) = \alpha\Big(\widehat{\boldsymbol{F}}_k(x) + \widehat{\boldsymbol{F}}^c_k(x)\Big),$$

a necessary and sufficient condition for which is that $\widehat{\boldsymbol{F}}_k(x)$ and $\alpha\widehat{\boldsymbol{F}}_N(x)$ have the same sign, which requires that

$$\text{sign}\Big(\widehat{\boldsymbol{F}}_N(x)\Big) = \text{sign}\Big(\widehat{\boldsymbol{F}}_k(x) + \widehat{\boldsymbol{F}}^c_k(x)\Big) = \text{sign}\Big(\widehat{\boldsymbol{F}}_k(x)\Big). \tag{S8}$$

Note that for $\alpha = 1$, it is trivially the case that the minimum of (S7) occurs for $k = N$, $\boldsymbol{\ell}(N,1) = 0$. This motivates taking $\alpha < 1 - \epsilon$ for some appropriate choice of $\epsilon$. Further, as $k \to N$ it is the case that the sign condition (S8) trivially becomes satisfied.

We define the *partial-sum relevance* at level $\alpha$ for a particular input $x \in \boldsymbol{\mathcal{B}}^+_{\tau_\ell;\mathsf{test}}$ as,

$$\mathsf{R}^\alpha_{\mathcal{D}}(x) = \text{argmin}_k\, \boldsymbol{\ell}(x;\alpha),\;\; \alpha \in (0, 1-\epsilon) \;\text{ subject to the constraint Eq. (S8)}, \tag{S9}$$

which provides an integer value $k^*(x) = \mathsf{R}^\alpha_{\mathcal{D}}(x)$ that gives the minimum number of active prototypes (the number of terms in the optimal partial-sum approximation $\widehat{\boldsymbol{F}}_{\boldsymbol{k}^*(x)}(x)$) needed to estimated $100\alpha$ percent of the total-sum approximation $\widehat{f}(x) = \widehat{\boldsymbol{F}}_N(x)$. Once $k^*(x)$ is at hand we can form a set of locally active prototypical examples that inform the classification decision for the vector $x$:

$$\mathsf{AES}(x) = \big\{\boldsymbol{i}_{y_i} \mid 1 \leq j \leq \boldsymbol{k}^*(x)\big\}. \tag{S10}$$

[1]Note that because $k$ is integer valued we have an integer programming problem. Also note that we can relate $\boldsymbol{\ell}(k;\alpha)$ to a simple quadratic loss function via $\widehat{f}^2(x)\boldsymbol{\ell}^2(k;\alpha) = \Big(\widehat{\boldsymbol{F}}_k(x) - \alpha\widehat{f}(x)\Big)^2$. Because the partial-sum approximation $\widehat{\boldsymbol{F}}_k(x)$ generally is not monotonic in $k$, minimizing absolute value loss generally gives a different answer than minimizing quadratic loss. Although one might prefer to replace the approximation $\widehat{f}(x)$ by the actual value $f(x)$ in (S7), we find that because the error between the two is generally small there is no practical harm in our choice, while the gain in insight and computation convenience when using (S7) is found to be useful.

[2]Alternatively to the use of (S7), one might opt to use the related regularized quadratic loss function given by (dropping the $x$-dependence in the notation)

$$\boldsymbol{\ell}'(k;\gamma) = \Big(\widehat{f} - \widehat{\boldsymbol{F}}_k\Big)^2 + \gamma\widehat{\boldsymbol{F}}^2_k \tag{S5}$$

$$\alpha = \frac{1}{1+\gamma} \in [0,1) \iff \gamma = \frac{\alpha}{1-\alpha} \in [0,\infty). \tag{S6}$$

Indeed, it is readily verified that

$$\boldsymbol{\ell}'(k;\gamma) = \frac{\widehat{f}^2}{\alpha}\,\boldsymbol{\ell}^2(k;\alpha) + (1-\alpha)\widehat{f}^2.$$

It is evident that $\alpha$ and $\gamma$ serve as closely related regularization parameters (and entirely equivalent regularization parameters in the case of quadratic loss); however we find the choice of the parameter $\alpha$ in the loss function (S7) to be more intuitively interpretable—see the subsequent discussion and, in particular, Footnote 20.

[3]In which case the value of the scalar, real-valued partial-sum threshold-function approximation $\widehat{\boldsymbol{F}}_k(x)$ "climbs up to" (or "explains") $100\alpha\%$ of the value of the scalar, real-valued total-sum threshold-function approximation $\widehat{\boldsymbol{F}}_N(x)$. (Recall that $x$ is classified as a mitotic figure when the transformed belief function $f(x) \approx \widehat{\boldsymbol{F}}_N(x)$ exceeds the transformed decision threshold $0 = \mathsf{SBC}(\tau_0;\tau_0)$—see Eq. (4).)

Note that this is equivalent to Eq. (13) when one sets $\mu_2 = |t_{\boldsymbol{k}^*(x)}|$. Further, it is evident that $|\mathsf{AES}(x)| = \boldsymbol{k}^*(x) = \mathsf{R}^{\alpha}_{\mathcal{D}}(x)$.

We then define the *sample mean statistic* Mean-RS *at percent level* $100\alpha\%$ by

$$\text{Mean-RS} = \text{Mean-RS}^{\alpha}_{\mathcal{D}} = \widehat{\mathsf{E}}_{\boldsymbol{\mathcal{B}}^{+}_{\tau_\ell;\mathsf{test}}}\big\{\mathsf{R}^{\alpha}_{\mathcal{D}}(x)\big\} = \frac{1}{|\boldsymbol{\mathcal{B}}^{+}_{\tau_\ell;\mathsf{test}}|}\sum_{x\,\in\,\boldsymbol{\mathcal{B}}^{+}_{\tau_\ell;\mathsf{test}}}\mathsf{R}^{\alpha}_{\mathcal{D}}(x), \tag{S11}$$

where $\widehat{\mathsf{E}}_{\boldsymbol{\mathcal{B}}^{+}_{\tau_\ell;\mathsf{test}}}\{\cdot\}$ represents a sample average over the test data set $\boldsymbol{\mathcal{B}}^{+}_{\tau_\ell;\mathsf{test}}$. Similarly, we define the *sample median statistic* Med-RS *at percent level* $100\alpha\%$ by

$$\text{Med-RS} = \text{Med-RS}^{\alpha}_{\mathcal{D}} = \mathsf{Median}_{x\in\boldsymbol{\mathcal{B}}^{+}_{\tau_\ell;\mathsf{test}}}\big\{\mathsf{R}^{\alpha}_{\mathcal{D}}(x)\big\}. \tag{S12}$$

We want these measures to be small, so that only a few number of informative prototype examples appear at the time of inference on a new datum $x$. It's worth noting that the Med-RS score has conceptual similarities to the median number of prototypes used for decision-making, as discussed in Arık and Pfister (2020).

Note that via the determination of the set, $\mathsf{AES}(x)$, of local prototypical examples we are conceptually replacing the approximation $\widehat{f}(x) = \widehat{\boldsymbol{F}}_N(x)$ of Eq. (S1) with the sparser, and generally more information-dense, approximation,[4]

$$f(x) \approx \widehat{f}^{*}_{\boldsymbol{k},\boldsymbol{c}}(x) \overset{\text{def}}{=} \widehat{\boldsymbol{F}}_{\boldsymbol{k}^*(x)}(x) = \sum_{\substack{i_{y_j}\,\in\,\mathsf{AES}(x)\\ j\,=\,1}}^{k^*(x)} \underbrace{c_{y_j}\,\rho^{(\rho_o)}_{k_{y_j},y_j}(x)}_{t_j(x)} \quad \text{where} \quad k^*(x) = |\mathsf{AES}(x)|. \tag{S13}$$

For a given truncation parameter $k$, we can form an objective measure of (local, $x$-dependent) predictive importance of $\widehat{\boldsymbol{F}}_{\boldsymbol{k}}(x)$ via the measure

$$\mathbf{PI}_1(k;x) = 1 - \frac{|f(x) - \widehat{\boldsymbol{F}}_{\boldsymbol{k}}(x)|}{|f(x)|}$$

or the measure[5]

$$\mathbf{PI}_2(k;x) = 1 - \frac{|\widehat{\boldsymbol{F}}_N(x) - \widehat{\boldsymbol{F}}_{\boldsymbol{k}}(x)|}{|\widehat{\boldsymbol{F}}_N(x)|} = 1 - \frac{|\widehat{\boldsymbol{F}}^{c}_{\boldsymbol{k}}(x)|}{|\widehat{\boldsymbol{F}}_N(x)|}.$$

From these we can form objective measures of $\mathsf{AES}(x)$ predictive importance via

$$\mathbf{PI}_1(k^*(x);x) = 1 - \frac{|f(x) - \widehat{f}^{*}_{\boldsymbol{k},\boldsymbol{c}}(x)|}{|f(x)|} = 1 - \frac{|f(x) - \widehat{\boldsymbol{F}}_{\boldsymbol{k}^*(x)}(x)|}{|f(x)|}$$

or

$$\mathbf{PI}_2(k^*(x);x) = 1 - \frac{|\widehat{f}(x) - \widehat{f}^{*}_{\boldsymbol{k},\boldsymbol{c}}(x)|}{|\widehat{f}(x)|} = 1 - \frac{|\widehat{\boldsymbol{F}}_N(x) - \widehat{\boldsymbol{F}}_{\boldsymbol{k}^*(x)}(x)|}{|\widehat{\boldsymbol{F}}_N(x)|} = 1 - \frac{|\widehat{\boldsymbol{F}}^{c}_{\boldsymbol{k}^*(x)}(x)|}{|\widehat{\boldsymbol{F}}_N(x)|}.$$

Because $\mathbf{PI}_1(\cdot;x)$ is baselined to the (transformed) belief function $f(x)$ learned by the Faster R-CNN, it is meaningful to compute $\mathbf{PI}_1(N,x)$ and thereby form a relative (relative to the total-sum predictor $\widehat{\boldsymbol{F}}_N(x)$) measure of predictive importance of a truncated sum predictor $\widehat{\boldsymbol{F}}_k(x)$ via

$$\mathbf{RI}(k;x) = \frac{\mathbf{PI}_1(k;x)}{\mathbf{PI}_1(N;x)}.$$

[4]Recall the ordering convention (S4).

[5]These measures are not guaranteed to be nonnegative.

In particular, one can form a measure of relative prediction performance of the $\mathsf{AES}(x)$ set as,

$$\mathbf{RI}(k^*(x);x) = \frac{\mathbf{PI}_1(k^*(x);x)}{\mathbf{PI}_1(N;x)}.$$

Given a test data set $\boldsymbol{\mathcal{B}}^+_{\tau_\ell;\mathsf{test}}$, we can form sample-averaged measures from the above local ($x$-dependent) measures in the obvious way:

$$\text{Mean-}\mathbf{PI}_1(k) = \widehat{\mathsf{E}}_{\boldsymbol{\mathcal{B}}^+_{\tau_\ell;\mathsf{test}}}\big\{\mathbf{PI}_1(k;x)\big\} = \frac{1}{|\boldsymbol{\mathcal{B}}^+_{\tau_\ell;\mathsf{test}}|}\sum_{x\,\in\,\boldsymbol{\mathcal{B}}^+_{\tau_\ell;\mathsf{test}}} \mathbf{PI}_1(k;x), \tag{S14}$$

$$\text{Mean-}\mathbf{PI}_2(k) = \widehat{\mathsf{E}}_{\boldsymbol{\mathcal{B}}^+_{\tau_\ell;\mathsf{test}}}\big\{\mathbf{PI}_2(k;x)\big\} = \frac{1}{|\boldsymbol{\mathcal{B}}^+_{\tau_\ell;\mathsf{test}}|}\sum_{x\,\in\,\boldsymbol{\mathcal{B}}^+_{\tau_\ell;\mathsf{test}}} \mathbf{PI}_2(k;x), \tag{S15}$$

$$\text{Mean-}\mathbf{RI}(k) = \widehat{\mathsf{E}}_{\boldsymbol{\mathcal{B}}^+_{\tau_\ell;\mathsf{test}}}\big\{\mathbf{RI}(k;x)\big\} = \frac{1}{|\boldsymbol{\mathcal{B}}^+_{\tau_\ell;\mathsf{test}}|}\sum_{x\,\in\,\boldsymbol{\mathcal{B}}^+_{\tau_\ell;\mathsf{test}}} \mathbf{RI}(k;x). \tag{S16}$$

In a similar manner, one can compute other sample statistical measures of performance, such as the median, etc. (see, for example, Eq. (S12)).

### S4 On concentration parameters and trimming conditions

Our RBF basis functions are trimmed (truncated to zero) to achieve further localization, according to the value of a trim parameter $\uprho_o$ in the following way:

$$\rho^{(\uprho_o)}_{k_y,y}(x) = \begin{cases} \rho_{k_y,y}(x), & \text{if } \rho_{k_y,y}(x) \geq \uprho_o \\ 0, & \text{otherwise} \end{cases}$$

for $0 \leq \uprho_o \leq 1$. Relations between the concentration parameters $k_y$ and trim parameter $\rho_0$ has been explored in the Supplementary section S4. Setting $\uprho_o = 1$ results in $\rho^{(1)}_{k_y,y}(x) = 0$ for all $x$ not directly proportional to $y$; i.e.,

$$\rho^{(1)}_{k_y,y}(x) = \mathbb{I}\{\hat{x} = \hat{y}\}\,.$$

For fixed $y$, non-trimming, $\rho^{(\uprho_o)}_{k_y,y}(x) = \rho_{k_y,y}(x)$, occurs for a particular value of $\uprho_o$ and $x$ when[6]

$$1 \geq \rho_{k_y,y}(x) \geq \uprho_o \geq 0 \iff \|\hat{r}_x - \hat{r}_y\|^2 \leq \min\left\{4, -\frac{2}{k_y}\ln\uprho_o\right\} = \min\left\{4, -2v_y\ln\uprho_o\right\}$$

which corresponds to the condition

$$\|\hat{r}_x - \hat{r}_y\| \leq \min\left\{2, s_y\sqrt{-2\ln\uprho_o}\right\} \tag{S17}$$

or, equivalently, when the following cosine similarity condition holds,

$$-1 \leq \max\left\{1 + v_y \ln\uprho_o, -1\right\} \leq \cos(\hat{r}_x, \hat{r}_y) \leq 1. \tag{S18}$$

In particular note that for $\uprho_0 = 0$ trimming is inactive for all $x$ and $y$,

$$\rho^{(0)}_{k_y,y}(x) = \rho_{k_y,y}(x),\ \forall x, y\,,$$

[6]Recall that $k_y = 1/v_y = 1/s_y^2$. Also, because $\hat{r}_x$ and $\hat{r}_y$ are unit vectors, we have $\|\hat{r}_x - \hat{r}_y\| \leq \|\hat{r}_x\| + \|\hat{r}_y\| = 2$ and $\|\hat{r}_x - \hat{r}_y\|^2 \leq 4$ where the upper bounds are attainable.

since

$$\|\hat{r}_x - \hat{r}_y\| \leq \|\hat{r}_x\| + \|\hat{r}_y\| = 2 = \lim_{\rho_0 \to 0} \min\left\{2, s_y\sqrt{-2\ln \rho_o}\right\}.$$

In our experiments we set the trim parameter to $\rho_0 = 0.1$, giving the non-trimming condition

$$\rho_0 = 0.1 \iff \|\hat{r}_x - \hat{r}_y\| \leq \min\{2,\, 2.15 s_y\}. \tag{S19}$$

Thus, when $2.15 s_y \leq 2$ then we have $\rho^{(0.1)}_{k_y,y}(x)$ is nonzero if $\hat{r}_x$ is within 2.15 $y$-scale units $s_y$ of $\hat{r}_y$ in the normalized representation space.

Note the $\rho_0$-dependent constraint that Eq. (S18) places on value of $k_y$ for the RBF to invoke the non-trimming condition $\rho^{(\rho_o)}_{k_y,y}(x) \neq 0$ in Eq. (6) for all $x$ and a *given*, fixed center representation vector $y$,

$$1 + v_y \ln \rho_o \geq -1 \iff k_y \geq -\frac{1}{2}\ln \rho_o. \tag{S20}$$

For example, the value $\rho_o = 0.1$ results in non-trimming for all $x$ for a given fixed $y$ if and only if

$$k_y \geq -\frac{1}{2}\ln \rho_o = -\frac{1}{2}\ln(0.1) = 1.15. \tag{S21}$$
